# Coherent chaos in a recurrent neural network with structured connectivity

**DOI:** 10.1101/350801

**Authors:** Itamar Daniel Landau, Haim Sompolinsky

## Abstract

We present a simple model for coherent, spatially correlated chaos in a recurrent neural network. Networks of randomly connected neurons exhibit chaotic fluctuations and have been studied as a model for capturing the temporal variability of cortical activity. The dynamics generated by such networks, however, are spatially uncorrelated and do not generate coherent fluctuations, which are commonly observed across spatial scales of the neocortex. In our model we introduce a structured component of connectivity, in addition to random connections, which effectively embeds a feedforward structure via unidirectional coupling between a pair of orthogonal modes. Local fluctuations driven by the random connectivity are summed by an output mode and drive coherent activity along an input mode. The orthogonality between input and output mode preserves chaotic fluctuations even as coherence develops. In the regime of weak structured connectivity we apply a perturbative approach to solve the dynamic mean-field equations, showing that in this regime coherent fluctuations are driven passively by the chaos of local residual fluctuations. Strikingly, the chaotic dynamics are not subdued even by very strong structured connectivity if we add a row balance constraint on the random connectivity. In this regime the system displays longer time-scales and switching-like activity reminiscent of “Up-Down” states observed in cortical circuits. The level of coherence grows with increasing strength of structured connectivity until the dynamics are almost entirely constrained to a single spatial mode. We describe how in this regime the model achieves intermittent self-tuned criticality in which the coherent component of the dynamics self-adjusts to yield periods of slow chaos. Furthermore, we show how the dynamics depend qualitatively on the particular realization of the connectivity matrix: a complex leading eigenvalue can yield coherent oscillatory chaos while a real leading eigenvalue can yield chaos with broken symmetry. We examine the effects of network-size scaling and show that these results are not finite-size effects. Finally, we show that in the regime of weak structured connectivity, coherent chaos emerges also for a generalized structured connectivity with multiple input-output modes.

**Author Summary:** Neural activity observed in the neocortex is temporally variable, displaying irregular temporal fluctuations at every accessible level of measurement. Furthermore, these temporal fluctuations are often found to be spatially correlated whether at the scale of local measurements such as membrane potentials and spikes, or global measurements such as EEG and fMRI. A thriving field of study has developed models of recurrent networks which intrinsically generate irregular temporal variability, the paradigmatic example being networks of randomly connected rate neurons which exhibit chaotic dynamics. These models have been examined analytically and numerically in great detail, yet until now the intrinsic variability generated by these networks have been spatially uncorrelated, yielding no large-scale coherent fluctuations. Here we present a simple model of a recurrent network of firing-rate neurons that intrinsically generates spatially correlated activity yielding coherent fluctuations across the entire network. The model incorporates random connections and adds a structured component of connectivity that sums network activity over a spatial “output” mode and projects it back to the network along an orthogonal “input” mode. We show that this form of structured connectivity is a general mechanism for producing coherent chaos.

## Introduction

Firing-rate fluctuations and irregular spiking are ubiquitous in the neocortex[28, 3]. Furthermore this temporal variability is often observed to be correlated across spatial scales ranging from local cortical circuits to the entire brain: in local cortical circuits both in membrane potential fluctuations [40] and on the level of spiking [27, 4, 20, 21], in the coherency measured in brain-wide EEG signals [19, 1], and in the global signal observed across all voxels in fMRI measurements [26, 14, 18].

A class of theoretical models has successfully accounted for temporal variability via internally generated chaotic dynamics of recurrent networks, whether through excitation-inhibition balance in spiking models[38, 23] or the more abstract models of rate chaos in randomly connected networks[30]. Yet a key emergent feature of these models is the decorrelation of neural activity such that the macroscopic, population activity remains nearly constant in time. Population-wide coherence or synchrony can be generated in a variety of ways for example by introducing spatial modes with self-excitation, but this comes at a cost of drowning out the chaotic fluctuations and yielding fixed points [35]. Indeed a major challenge to theorists has been to produce network models which generate spatially coherent, temporally irregular fluctuations which can account for broad spatial correlations observed in experiments.

Two recent studies have shown that excitation-inhibition balance networks can generate spatially modulated correlations [25, 6]. In both of these studies the correlations are driven by common input from an external source, and the average correlation across the network remains small. It remains an open question whether a network can internally generate correlated fluctuations that are coherent across the entire network.

The chaotic dynamics of a network of randomly connected firing-rate neurons has been well-studied [30, 12]. In such a network each individual neuron’s firing rate is given by a non-linear function of its input, which is in turn a weighted sum of the firing rates of all other neurons in the network. The network exhibits a phase transition from a fixed point to chaotic activity in which the randomness of the weights reverberates uncorrelated fluctuations throughout the network. Typically in this chaotic regime pairwise correlations are small and no coherent fluctuations emerge. Here we extend this model by adding a low-rank structured component of connectivity to the random connections. The structured connectivity sums the fluctuations along one spatial mode and projects them along a second, orthogonal mode yielding coherent fluctuations without drowning out the individual neuron variability which continues to drive the chaotic dynamics. A previous work studied a specific example of this structure and focused primarily on analyzing the non-chaotic regime[8]. Here we focus on the chaotic regime and show that this form of structured connectivity together with random connections provides a basic mechanism for internally generating coherent fluctuations.

## Results

We study a network of *N* neurons in which the connectivity between neurons has two components: a random component, **J**, and a rank-1 structured component, 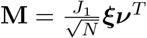, an outer product of a pair of orthogonal vectors both of which have elements of *O* (1) and norm 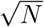, with strength parameter *J*_1_. We restrict the elements of ***ξ*** to be binary, *ξ_i_* = ±1, which will be important for some of the results to come, and we will comment on when this restriction can be relaxed. We can think of the row vector, ***ν**^T^*, as an “output mode” performing a read-out of the network activity, and the column vector, ***ξ***, as a corresponding “input mode” along which the output mode activity is fed back to the network (Fig 1A).

**Figure 1.**
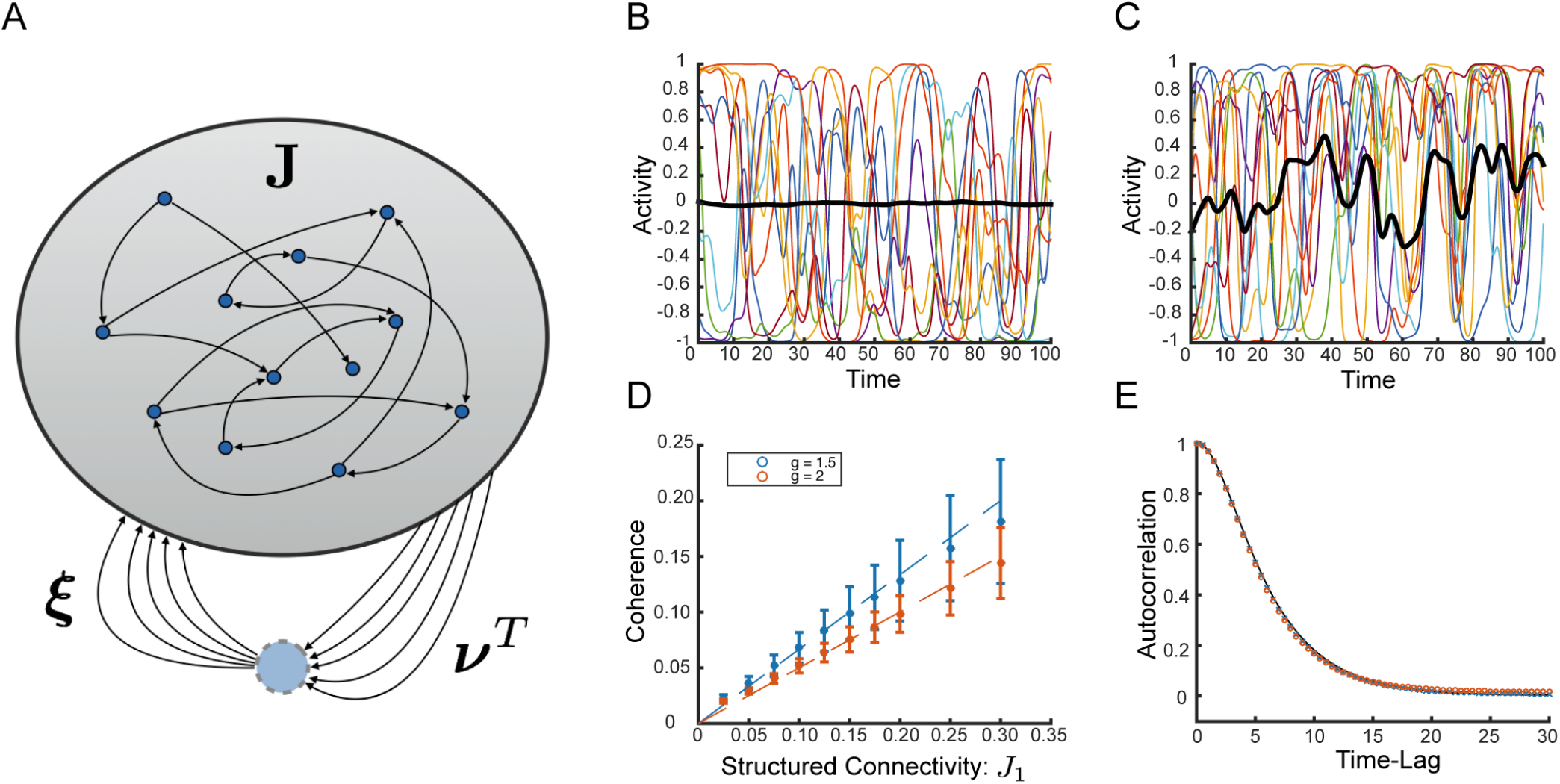
Random Network with Structured Connectivity Generates Coherent Chaos. **(A)** Network schematic showing neurons connected via random matrix **J** and rank-one structured connectivity. The structured component is represented schematically as a feed-forward loop with drive through the output mode, ***ν***, and feedback through the input mode, ***ξ***. In our model these two vectors are orthogonal. The standard deviation of the random component is given by 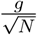 and the strength of the structured component is 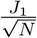. **(B)** Sample network dynamics without structured component, i.e. *J*_1_ = 0, colored traces show a random collection of ten neural activity patterns, *ϕ_j_*, black trace shows the coherent mode activity, 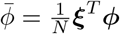. **(C)** Sample dynamics for *J*_1_ = 1. Coherent mode displays substantial fluctuations. **(D)** Coherence, *χ* (definition in text), as a function of the strength of structured connectivity component, *J*_1_ for small values of *J*_1_. Simulation and theory (valid in the weak structured connectivity regime - *J*_1_ ≪ *g*) shown for both *g* = 1.5 and *g* = 2. Bars show standard deviation over 60 realization of the random connectivity. **(E)** Passive coherent chaos. With weak structured connectivity fluctuations of the coherent mode follow the fluctuations of the independent residual components. Normalized autocorrelation of the coherent component of the current, *q*(*τ*), in red circles. Average normalized autocorrelation of the residuals, *q_δ_* (*τ*), in blue ‘x’s. Both are averaged over 60 realizations of the random connectivity with *J*_1_ = 0.1. Prediction from theory in black. *N* = 4000 and *g* = 2 in all panels unless stated otherwise.

The random component of the connectivity is given by the matrix **J** consisting of identically distributed independent Gaussian elements with mean 0 and variance 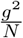, where *g* is an *O* (1) parameter.

The state of each neuron is defined by its synaptic current, *h_i_* (*t*), with its firing rate given by *ϕ_i_ ≡ ϕ* (*h_i_* (*t*)), where *ϕ* is a sigmoidal function. For some later results it will be necessary to assume that *ϕ* has a first derivative that is an even function. We therefore assume here for concreteness *ϕ* (*h*) = tanh (*h*) unless otherwise noted, and we comment on when this assumption can be relaxed.

The dynamics of the synaptic current vector, **h**, is given by

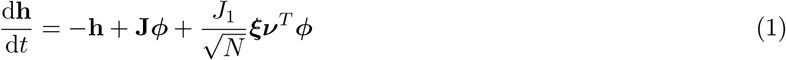

We have scaled the strength of the structured connectivity such that when the scaling-parameter *J*_1_ ∼ *O* (1) the contribution of the structure to individual synapses is of the same order of magnitude as the typical random connection.

We will be particularly interested in the coherent activity and the coherent current, i.e. the spatial overlap of both the firing rate and the synaptic current with the input mode. These are defined respectively as

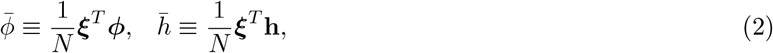

We define the measure of spatial coherence as the fraction of the total power of the network current *h_i_* that is shared along the input mode, ***ξ***:

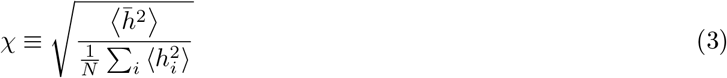

where 〈 〉 represents average over time. This is a useful measure as it varies from 0 to 1, and will yield 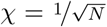 for entirely independent, uncoupled fluctuations, and *χ* = 1 for complete synchrony along ***ξ***.

Without the structured component, i.e. *J*_1_ = 0, the network exhibits a phase transition at *g* = 1 from a zero fixed point to chaos[30]. In the chaotic state the randomness of the connectivity decorrelates the input current and yields an asynchronous state in which neurons fluctuate with negligible correlations such that both *ϕ* and *h* are nearly constant in time, and 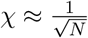 (Fig 1B).

Setting *J*_1_ = 1, we observe significant correlations along the input mode, such that the coherent mode activity, *ϕ*, fluctuates significantly. For this network with *N* = 4000 we find the coherence *χ* ≈ 0.4, which is about 25 times larger than is observed in the asynchronous state. (Fig 1C).

In order to analyze this system we decompose the dynamics of the synaptic currents into coherent component, *h*, and the vector of residuals currents, *δ***h** = **h** − h***ξ***. We also decompose the activity into its coherent component, *ϕ*, and vector of residual activity, *δ**ϕ*** = ***ϕ*** − *ϕ****ξ***. Because of the orthogonality between input and output mode, the output mode ignores the coherent component, *ϕ*, and projects only the activity of the residuals: ***ν**^T^ **ϕ*** = ***ν**^T^δ**ϕ***.

By averaging the dynamic equations 1 over the input mode, ***ξ***, we can write decomposed dynamics for the residual synaptic current vector and for the coherent current:

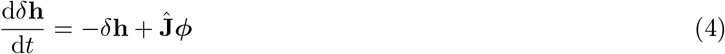

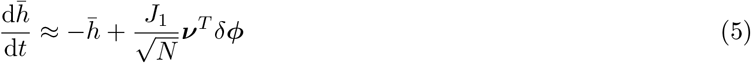

where the effective connectivity matrix in the residual dynamics (discussed more fully in Methods) is

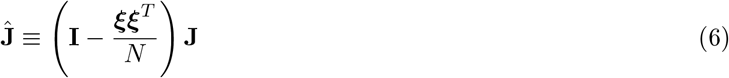

which projects the output of **J** into the (*N* − 1) dimensional subspace orthogonal to ***ξ***. This guarantees that the decomposed dynamics satisfy the constraint ***ξ**^T^ δ***h** = 0. For most of what follows this constraint can be ignored as it contributes only 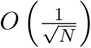 to each synaptic current. Nevertheless it plays a role later and we introduce it here for completeness. We have, however, ignored the small projection of the random connectivity along the coherent mode 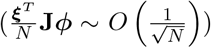. Note that even in this approximation, the nonlinearity of *ϕ* in these equations couples the coherent and residual degrees of freedom.

In order to attempt to solve the approximate system we could assume that *h* (*t*) fluctuates according to some known random process and then consider the dynamics of the *δh_i_* with the firing rate of individual units as given by *ϕ* (*δh_i_* (*t*) + *ξ_i_h* (*t*)). However, for general *J*_1_ we are unable to analytically close the loop and self-consistently compute the statistics of *h* (*t*).

### Weak Structured Connectivity Yields Passive Coherent Chaos

In order to proceed analytically, we take a perturbative approach, assuming *J*_1_ ≪ 1. In this regime we assume the fluctuations in *h* (*t*) are Gaussian and we turn to computing the autocorrelations of both the coherent component

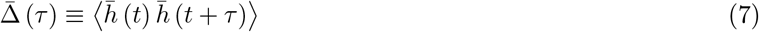

and of the residuals,

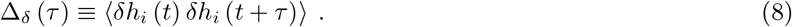

For small *J*_1_ we assume that the coherent current is small (*h* ≪ 1) and therefore in the dynamics of the residual currents (Eqn 4) we approximate 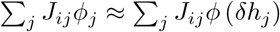. The result is that to leading order the autocorrelation of the residuals is given by the zeroth-order (*J*_1_ = 0) autocorrelation. That is, the residual currents fluctuate as independent Gaussian processes almost identically to the situation without structured connectivity. These residual fluctuations are summed over the output mode yielding substantial fluctuations in ***ν**^T^ δ**ϕ*** (recall, *ν_i_* ∼ *O* (1)). These in turn drive Gaussian fluctuations in the coherent mode and we show in Methods that its autocorrelation is given to first-order by

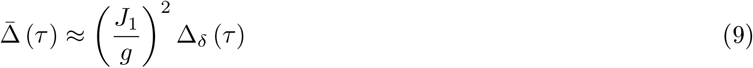

That is, to leading order the autocorrelation of the coherent component is simply a scaled version of the local, residual autocorrelation.

We verify this prediction numerically in Fig 1E for *J*_1_ = 0.1 showing the normalized autocorrelations, 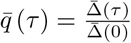 and *q_δ_* (*τ*) = 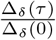 as well as the prediction from theory. Qualitatively this means that in this regime chaos is driven by the emergent fluctuations in the local synaptic current similar to in the *J*_1_ = 0 case, and that the coherent component can be said to absorb these fluctuations passively along the input mode, ***ξ***.

In this regime then, the coherence is given simply by

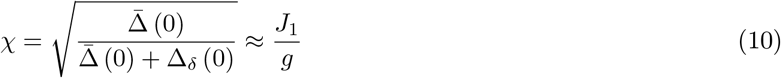

Numerically, we find that this approximation provides a good description of the system’s state for up to *^J^*_1_/*_g_* ≈ 0.2 (Fig 1D).

We can understand the coherent chaos in this regime qualitatively: the residual synaptic currents driven by the random connectivity fluctuate as uncorrelated Gaussian processes, and the resulting independent fluctuations in firing rates will be summed over the output mode, ***ν***, which projects to the input mode, ***ξ***, driving coherent fluctuations in *h* (*t*). If the two modes, ***ξ*** and ***ν*** had substantial overlap then the coherent fluctuations along ***ξ*** would drive positive feedback through ***ν*** driving the neurons to a fixed point. The orthogonality of these modes effectively embeds a feedforward structure from the output mode, ***ν***, to the input mode, ***ξ***, within the recurrent connectivity. This enables the persistence of stable fluctuations along the input mode, ***ξ***, which do not feedback to ***ν***, thus preventing either saturation or oscillations.

Importantly, in the regime of passive coherence we can relax the restrictions on ***ξ*** and *ϕ*: Our results here hold for any smooth, sigmoidal non-linearity and for any ***ξ*** which is norm 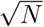 and orthogonal to ***ν***. In fact approximate orthogonality is sufficient in this regime: structured connectivity consisting of an outer product of two randomly chosen vectors will generate mildly coherent fluctuations.

### Random Connectivity with “Row Balance”

We observe that as *J*_1_ increases to values of *O* (1), the network displays significant variability in the dynamics from realization to realization. The coherent mode autocorrelation function, Δ (*τ*), for example, is no longer self-averaging (Fig 2A). As we increase system size, *N*, we find that the realization-to-realization variability in Δ (0) saturates to a finite value (Fig 2B). Moreover, when *J*_1_ ≳ 1 we observe realization-dependent transitions out of chaos to either fixed points or limit cycles (Fig 2C).

**Figure 2.**
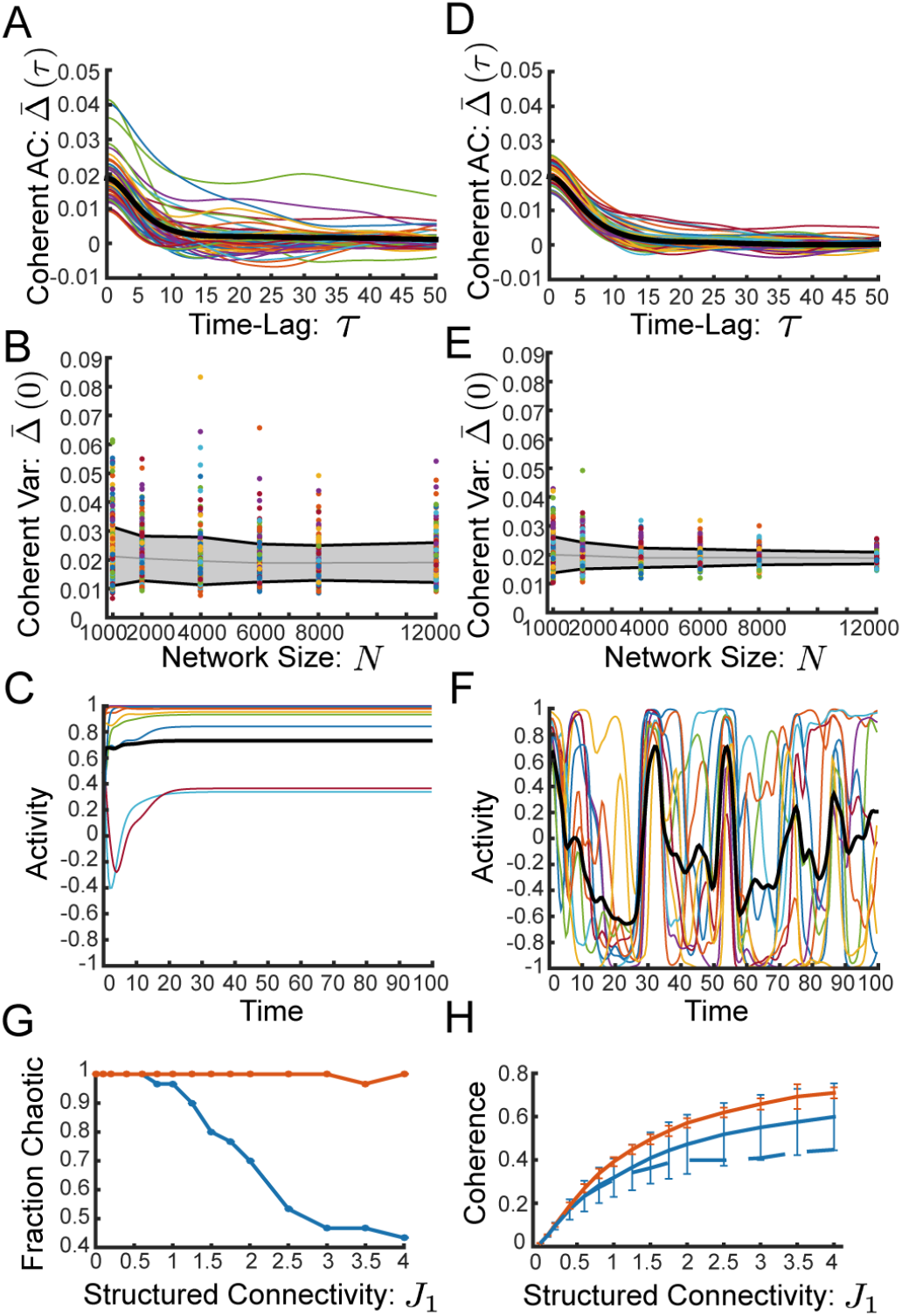
Row Balance Preserves Chaos, and Increases Coherence. **(A)** Full coherent mode autocorrelation, Δ (*τ*) of 60 individual realizations. Thick black line shows average over realizations. Network with random connectivity **J** without row balance exhibits significant difference between realizations **(B)** Coherent mode variance, Δ (0), as a function of network size for 300 individual realizations. Gray line shows average and gray region with black boundary shows one standard deviation over realizations. Without row balance the standard deviation (over realizations) saturates to a finite value as the network size increases, indicating that the variability between realizations is not a finite-size effect. **(C)** Without row balance a network with moderate structured connectivity (*J*_1_ = 2.5) exhibits a fixed point. **(D)-(F)** Same as (A)-(C) respectively, but network has “row balance” random connectivity, 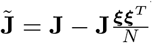. **(D)** Individual realizations of Δ (*τ*) are all very close to the average. **(E)** With row balance the standard deviation of Δ (0) over realizations shrinks with *N*, suggesting that the variability between realizations is a finite-size effect. **(F)** Same realization of **J** as in (C), but with row balance. Chaos is preserved. **(G)-(H)** Networks without row balance in blue, with row balance in red **(G)** Fraction of realizations (out of 30 realizations) that lead to chaotic dynamics, as a function of structural connectivity, *J*_1_. Row balance keeps nearly all realizations chaotic. **(H)** Coherence, *χ*, as a function of *J*_1_computed for the realizations from (E). Row balance increases coherence. Error bars display standard deviation. Full line shows all realizations, dashed line displays average coherence restricted to the chaotic realizations. *J*_1_ = 0.2, *g* = 2, and *N* = 4000 for all panels unless otherwise noted.

The reason for this realization dependence is that as *J*_1_ increases and the fluctuations in the coherent mode grow, feedback is generated through the interaction between the random connectivity, **J**, and the input and output modes, ***ξ*** and ***ν***. First of all, **J** maps the coherent activity back along the input mode with a small realization-dependent component which we ignored in Eqn 5 driving feedback directly to the coherent current, *h*. Secondly, **J** maps the coherent activity along the the residuals in a realization-dependent direction biasing the residual fluctuations *δ***h**. This direction in turn will have realization-dependent component along the output mode, ***ν***, and therefore the coherent activity may additionally drive feedback by pushing the residual fluctuations along the output mode. See Methods for more details.

To suppress the strong realization dependence of the dynamics, we refine the random connectivity matrix by defining

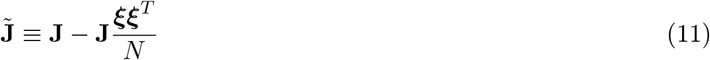

This subtracts from each row of **J** its weighted average along the input mode. This “row balance” subtraction has been previously observed to remove realization-dependent outliers from the eigenspectrum of the full connectivity matrix [22, 34].

We find here that row balance suppresses realization-to-realization variability in the nonlinear chaotic dynamics, for example reducing the variability in the autocorrelation of the coherent mode, Δ (*τ*) (Fig 2D). We observe that with row balance this variability drops as a function of increasing system size, suggesting (although not proving) that the dynamics are now self-averaging in the limit of large *N*, at least for these values of *J*_1_ (Fig 2E).

The impact of row balance on the chaotic dynamics can be understood by noting that the resulting connectivity matrix, **J**̃, now has a null-space, and the input mode, ***ξ***, lies within it (**J**̃***ξ*** = 0). The result of row balance then is to ensure that the random connectivity matrix filters out any coherent activity fluctuations, *ϕ* (*t*):

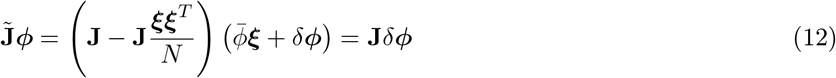

This prevents the coherent mode activity from driving feedback to the dynamics of the coherent current (Methods).

Interestingly, we find that row balance allows chaotic fluctuations to persist for larger values of *J*_1_, whereas without row balance the majority of realizations exhibit fixed points or limit cycles for *J*_1_ ≳ 3 (Fig 2F-G). Furthermore, the chaotic dynamics are more coherent with row balance than without (Fig 2H). As the structured connectivity is made stronger row balance appears to enable the dynamics to grow increasingly coherent even as chaotic fluctuations persist.

### Row Balance Yields Self-Tuned Coherent Chaos

When we increase the strength of structured connectivity, *J*_1_, we find that with row balance the network yields chaotic dynamics which are strikingly coherent and display switching-like macroscopic activity (Fig 3A). In contrast to the case of weak structured connectivity, the coherent mode dynamics are no longer passively driven by the fluctuations in the residual synaptic currents. This is evidenced by the normalized autocorrelation of the coherent mode, *q*(*τ*), which is no longer close to the normalized residual autocorrelation, *q_δ_* (*τ*), but rather has qualitatively different dynamics including longer time-scales (Fig 3B as compared to Fig 1E).

**Figure 3.**
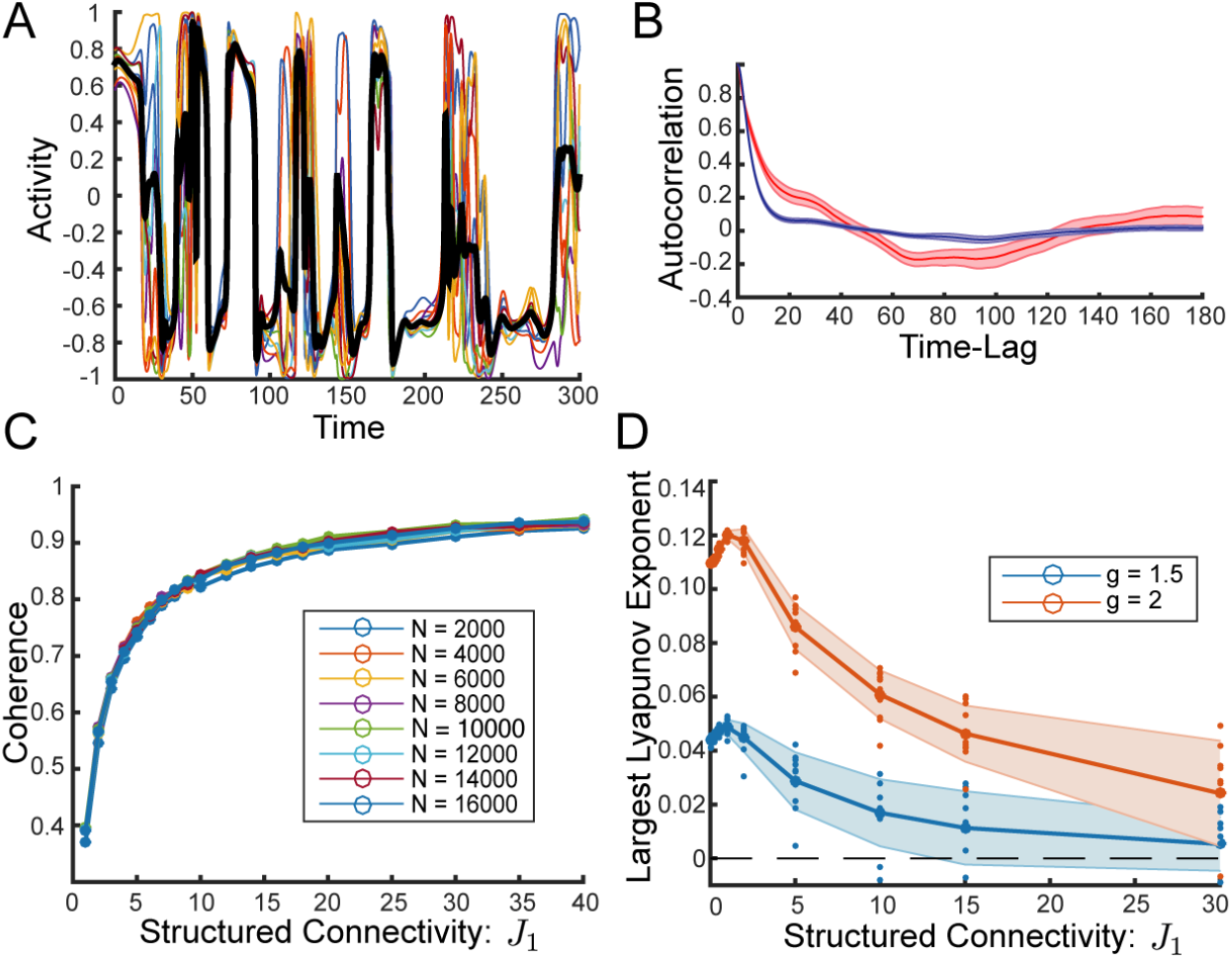
Strong Structured Connectivity with Row Balance Generates High Coherence Even as Chaos Persists. **(A)** Sample activity of 10 randomly chosen neurons, *ϕ_j_*, and coherent mode activity, *ϕ*, in black. Strong structured connectivity with row balance subtraction to the random component of connectivity yields chaotic activity that is highly coherent with switching-like behavior. **(B)** Normalized autocorrelation of coherent mode, *q*(*τ*), in red. Average normalized autocorrelation of the residuals, *q_δ_* (*τ*), in blue. Shaded regions show standard deviation over 25 initial conditions of the same connectivity. Strong structured connectivity yields coherent mode dynamics that are qualitatively different from those of the residuals. *J*_1_ = 15.8 for both (A) and (B), compare to Fig 1C and Fig 1E, respectively. **(C)** Coherence, *χ*, as a function of *J*_1_ is independent of network size. Coherence appears to approach 1 as *J*_1_ is increased demonstrating that chaos persists even as fluctuations in the residuals shrink (See also Fig S4). Coherence is averaged over 30 realizations of the connectivity for each *N*, excluding the few fixed points and limit cycles that occur for larger *J*_1_ (2 out of 30 or less for the largest values of *N*). **(D)** Largest Lyapunov exponent as a function of *J*_1_. Thick line shows average over 10 realizations, small dots show values for individual realizations, and shaded region is standard deviation. All but a small fraction of realizations are chaotic, even in the region where *χ >* 0.9. *N* = 4000 and *g* = 2 in all panels unless noted otherwise.

We find that the coherence, *χ*, increases steadily as a function of the strength of structured connectivity, *J*_1_, and notably it is independent of system size. (Fig 3C).

To check whether this highly coherent state is chaotic, we calculate the largest Lyapunov exponent and verify that the dynamics are indeed chaotic for a vast majority of realizations even as the fluctuations are highly coherent (Fig 3D).

We now examine the qualitative changes in the chaotic state of the network with row balance as *J*_1_ increases. We observe that for *J*_1_ ≲ 1 the fluctuations are unimodal with an approximate Gaussian shape. The temporal fluctuations are dominated by a single time constant as with zero *J*_1_ (Fig 4A-B). On the other hand, for larger values of *J*_1_ the fluctuations deviate dramatically from Gaussian and instead become sharply bimodal (Fig 4A).

**Figure 4.**
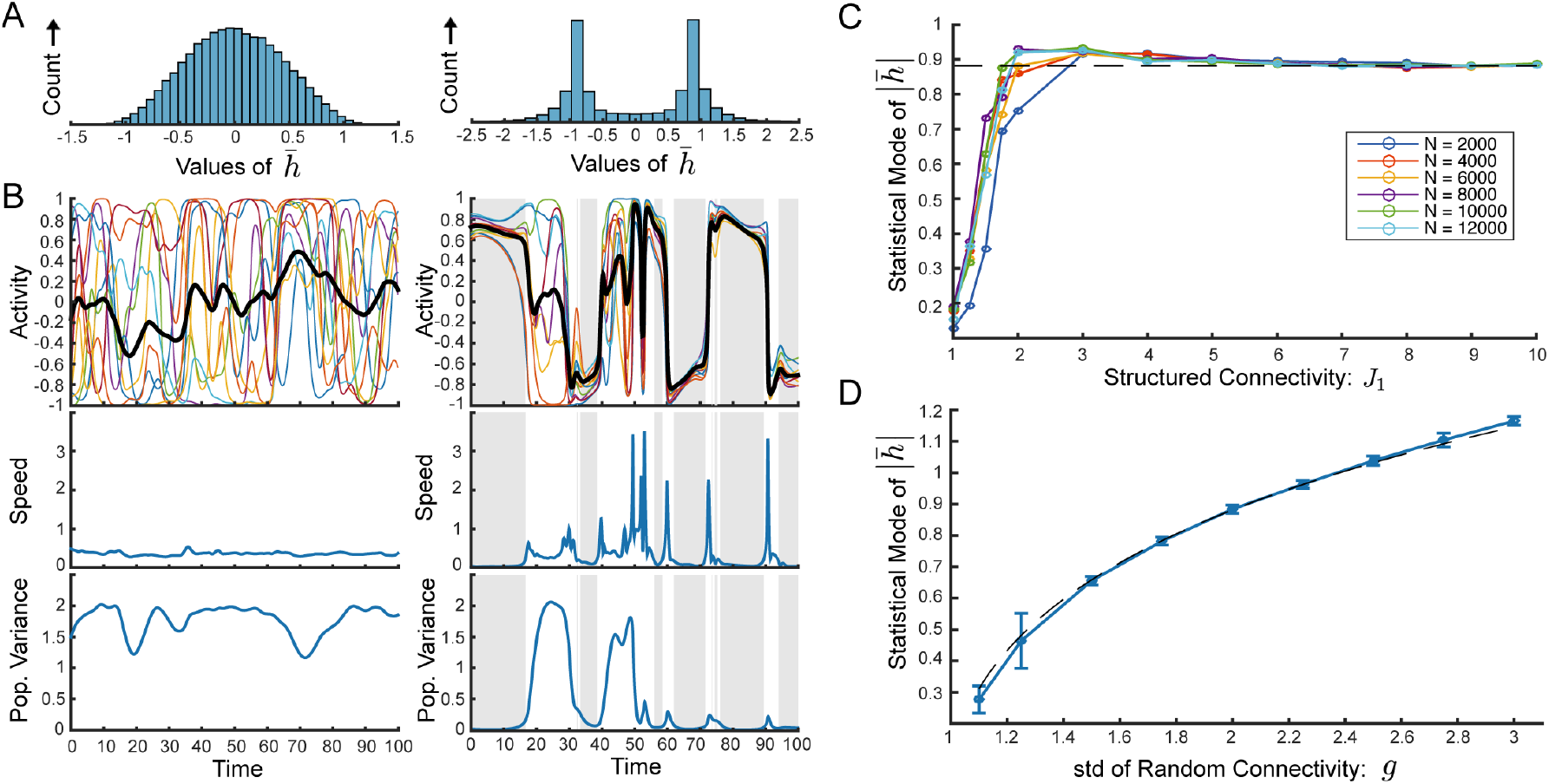
Self-Tuned Coherent Chaos. **(A)-(B)** Comparison between weak structured connectivity (Left: *J*_1_ = 0.8) and stronger structured connectivity (Right: *J*_1_ = 15.8). Both with *N* = 4000. **(A)** Histogram of values of the coherent mode current, *h* (*t*). Mild structured connectivity yields coherent fluctuations with a peak at zero and a distribution that appears not far from Gaussian. For stronger structured connectivity the histogram is clearly non-Gaussian and highly bimodal. **(B)** Top: Sample activity of 10 randomly chosen neurons, *ϕ_i_* (*t*) and coherent mode activity, *ϕ* (*t*). Middle: Speed of network during same epoch of activity, defined as the per-neuron norm of the rate-of-change vector, i.e. 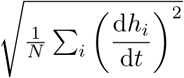. Bottom: Instantaneous population variance of the residual currents *δh_i_* (*t*). For mild structured connectivity, *ϕ* (*t*) fluctuates around zero (top), speed is roughly constant throughout the trial (middle), residual currents maintain large variance throughout (bottom). On the other hand, for stronger structured connectivity, there is state-switching between bouts of high and low coherent-mode activity (top), these same bouts are associated with vanishing speed (middle), and with small residual currents (bottom). **(C)** Statistical mode (most frequent value) of |*h*| as a function of *J*_1_. The results indicate a crossover to self-tuned coherent chaos, defined by the bimodal peaks of |*h*| reaching a constant value. The crossover occurs very rapidly and independently of *N*. Dashed line shows 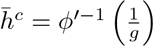 **(D)** The statistical mode of |*h*| as a function of *g* with fixed *J*_1_ = 10 and *N* = 8000. Dashed line shows 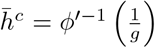. *g* = 2 in all other panels.

Furthermore, we find that with larger values of *J*_1_ the network exhibits intermittent switching between two different values of *h* (Fig 4B, top right). We observe that the dynamics at both of these values of *h* are qualitatively distinct as reflected by the speed of the dynamics and the level of coherence. We define the speed of the dynamics as the norm of the vector of first time-derivatives per neuron: 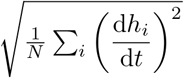. Similar measures have been used to find fixed points and slow dynamics in highly non-linear dynamics [32]. We observe that the two distinct values of *h* are both associated with vanishing speed (Fig 4B, middle right). Additionally they are associated with very high levels of coherence, i.e. small residuals as quantified by the population variance, 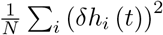 (Fig 4B, bottom right). Evidently, in the network with larger values of *J*_1_ there are two distinct states with slow dynamics and high levels of coherence, and the network switches rapidly between these two. In contrast, with mild structured connectivity both the speed and the variance of the residuals remain roughly constant throughout the trial (Fig 4B, left).

To gain insight into the emergence of the switching dynamics, we examine the most frequent value of |*h* (*t*)| as a function of *J*_1_ and we find that there is a rapid crossover from a state with unimodal fluctuations around zero, to a state with bimodal peaks and that the most frequent value of |*h* (*t*)| saturates quickly with *J*_1_ and then remains constant (Fig. 4C). What is the nature of this regime? And what determines the two values of *h* that come to dominate the dynamics?

Because the bouts of slow dynamics are associated with small residuals, we assume *δh_i_* ≪ 1 and approximate ***ϕ*** (*t*) ≈ *ϕ* (*t*) ***ξ*** + *ϕ′*(*h* (*t*)) *δ***h**. Note that we have made use of the fact that *ξ_i_* = ±1 and that *ϕ* is an odd function. Because ***ξ*** is in the null-space of the row-balanced connectivity (**J̃***ϕ* =**J***δ**ϕ***, as discussed above) the residual dynamics (Eqn 4) become

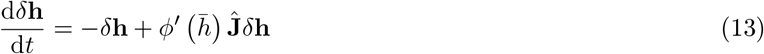

We observe that in these linearized dynamics the coherent mode current, *h* (*t*), plays the role of a dynamic gain via the slope of the transfer function, *ϕ′*.

In the linearized dynamics we turn to the eigenvectors, **u**^(*k*)^, of **Ĵ** and decompose the residual dynamics according to 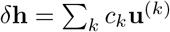. Given an instantaneous value of *h*, the independent dynamics of the eigenmodes are

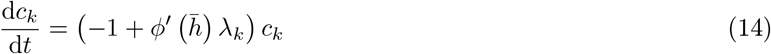

where *λ_k_* is the *k*th eigenvalue of **Ĵ**. (We show in Methods that **Jˆ** has the same eigenvalues as **J̃**)

The leading eigenvalue, *λ*_1_, has real part approximately equal to *g*. If *h* is small then *ϕ′*(*h*) ≈ 1 and there are many modes that diverge exponentially. If *h* is large (either positive or negative) then *ϕ′*(*h*) ≈ 0 and then all the modes decay exponentially. However, there are two critical values of *h* which yield marginal and therefore slow dynamics for the leading mode, *c*_1_. These are the values, *h*^*c*^, for which the slope of the transfer function is equal to 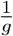:

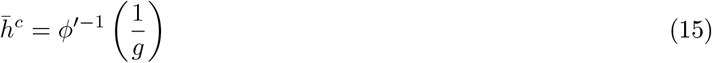

If *h* (*t*) ≈ *h*^*c*^ and the residuals are small then the time constant of fluctuations in the leading eigenmode, (1 − *ϕ′* (*h*) *g*)^−1^, are very long.

Indeed we find that the most frequent value of |*h* (*t*)| as a function of *g* fits the curve *h^c^* (*g*) very well (Fig. 4D).

We conclude that the switching between two states each with slow dynamics and a high level of coherence observed in Fig 4B reflects a regime of self-tuned coherent chaos in which the coherent mode self-adjusts to a critical value so that the dynamics of the small residuals are near-marginal, giving rise to slow dynamics. The above linearized dynamics (Eqn 13) are not exact and therefore non-linear interactions eventually destabilize the system and precipitate a state-switch. Nevertheless, the linearized dynamics dominate the dynamics of the small residuals during the bouts of high coherence.

As observed in Fig 4C, when we increase *J*_1_, the most frequent value of |*h* (*t*)| rapidly increases until it saturates at a value very near to 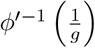. Moreover, we find that this crossover to the regime of self-tuned coherent chaos occurs at moderate values of *J*_1_ (on the order of *g*), independently of network size.

Notice the crucial role of row balance in facilitating self-tuned coherent chaos: row balance filters out the direct contribution of the coherent mode activity to the dynamics of the residuals and enables the coherent mode to act as a dynamic gain. The coherent mode then self-adjusts to cancel the leading eigenvalue of **Ĵ** and yield bouts of slow, highly coherent dynamics.

We note that we can loosen the constraint on the symmetry of the transfer function and allow any smooth sigmoidal transfer function if we restrict the input mode to be uniform, *ξ_i_* = 1 for all *i*. We show an example of the self-tuned coherent chaotic state for a non-symmetric transfer function in Fig S1.

### Symmetry breaking in the self-tuned chaotic regime and transition to fixed point

The example of Fig. 4A illustrates dynamics which reside in the positive and negative coherent states with equal frequency, maintaining the *h_i_* → *−h_i_* symmetry of the underlying dynamic equations. We observe that in may realizations this symmetry is violated at the single trial level for sufficiently strong *J*_1_, as demonstrated in Fig. 5A We measure the asymmetry in a single trial as the absolute value of the *time-averaged* coherent activity, |〈*h*〉|, and we find that asymmetry grows gradually with *J*_1_ throughout the chaotic regime, and at different rates for different realizations (Fig 5B).

**Figure 5.**
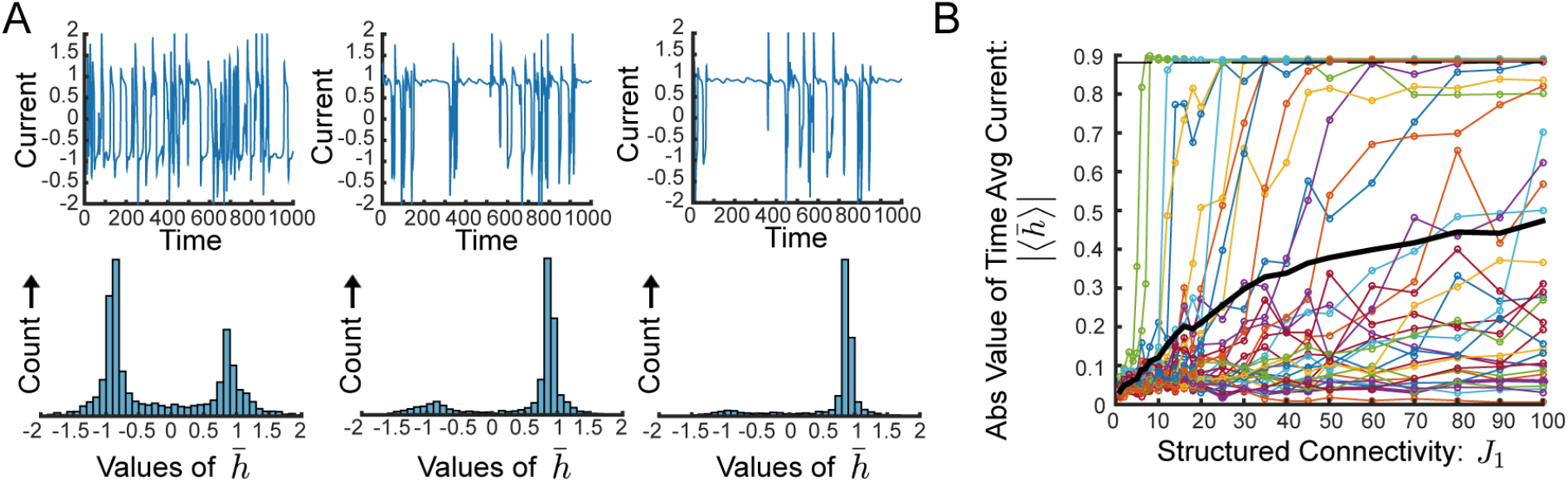
Realization-Dependent Symmetry Breaking in the Self-Tuned Chaotic Regime. **(A)** Sample traces of coherent mode current, *h* (*t*), (top) and histogram of values of *h* (*t*) (bottom) from a connectivity realization with real eigenvalue for *J*_1_ = 40, 60, 80 increasing from left to right. Dynamics exhibit pronounced asymmetry. **(B)** Absolute value of the time-average coherent mode current, |〈*h*〉|, as a function of *J*_1_. Each colored line represents a single connectivity realization, averaged over 10 initial conditions. For many individual realizations, |〈*h*〉| is significantly non-zero over a large range of values of *J*_1_, while still not arriving at fixed point value (displayed by dashed line). We display the 37 realizations with real leading eigenvalue out of 100 total realizations from this set of trials. Thick black line shows average over those realizations. *N* = 8000 for all panels.

As is evident in Fig. 5B, for many realization the level of asymmetry increases with *J*_1_ until it reaches a maximal value of 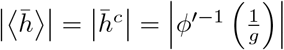 (dashed line). At this point, the system spends all the time at one of the possible states, suggesting a transition to fixed point.

Indeed, as has been previously reported by Garcia Del Molino et al [8], realizations of **Ĵ** that have a real leading eigenvalue (see also Methods) yield a fixed point equation for the above linearized dynamics (Eqns 13):

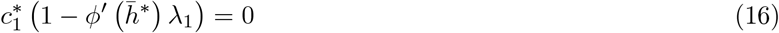

The fixed point requires 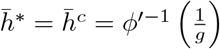 so that all but the leading eigenmode decay to zero, and the resulting fixed point for 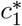 is marginally stable.

Indeed, we find that realizations of **Ĵ** with a real leading eigenvalue undergo a transition to a fixed point upon sufficient increase of *J*_1_. Similar to the preceding chaotic state, the fixed point is highly coherent, with very small residuals. Furthermore, it exhibits the hallmarks of the self-tuned coherent state: the value of the coherent mode is close to 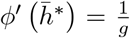, the residuals are aligned with the leading eigenvector of **Ĵ**, and the the leading eigenvalue of the Jacobian matrix at the fixed point is close to zero (Fig S2).

### Oscillatory fluctuations in self-tuned chaos and transition to limit cycle

The above symmetry breaking and transition to fixed point is observed only for some of the realizations of **J**. In most of the other cases, rather than symmetry breaking, we observe an increased oscillatory component in the chaotic dynamics. This is reflected in the presence of a large second peak in the normalized autocorrelation function, *q*(*τ*) (Fig 6A).

**Figure 6.**
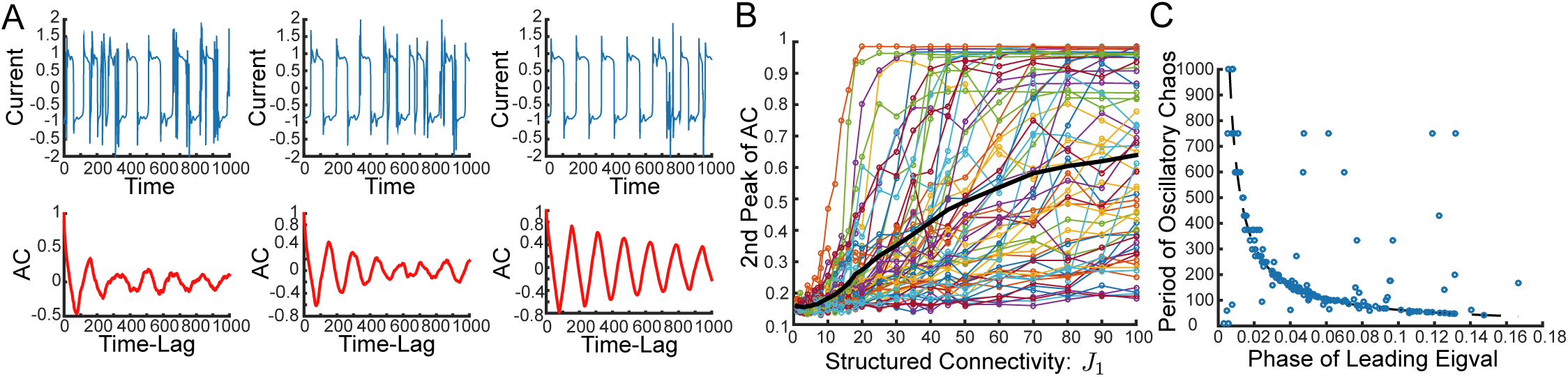
Realization-Dependent Oscillatory Imprint on the Self-Tuned Chaotic Regime. **(A)** Sample traces *h* (*t*) (top), and normalized autocorrelation *q*(*τ*) (bottom) of coherent mode current for a connectivity realization with complex eigenvalue for *J*_1_ = 25, 30, 35 increasing from left to right. Dynamics exhibit pronounced oscillatory power and the autocorrelation exhibits a pronounced peak near the same frequency that will dominate the limit cycle for larger *J*_1_. **(B)** Height of second peak of the autocorrelation of the coherent mode input as a function of *J*_1_. Each colored line represents a single connectivity realization, averaged over 10 initial conditions. For many realizations, there is a significant second peak in the autocorrelation over a long range of values of *J*_1_ well before a limit cycle is reached. We display the 63 realizations which had complex leading eigenvalue out of 100 in this set of trials. Thick black line shows average over those realizations. **(C)** Observed period of oscillatory chaos vs phase of leading eigenvalue for 181 realizations from which we were able to measure an oscillatory period with chaotic fluctuations (out of 196 realizations with complex leading eigenvalue in this set of trials. In order to confine to realizations and values of *J*_1_ that yielded chaos, we restrict to those with second peak of autocorrelation less than 0.8. These had average height of second peak over all realizations: 0.5). Dotted line shows prediction from theory: 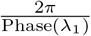. The bulk of realizations are very well predicted although a notable fraction are not. The median error of prediction was 7.75 (average period over these realizations: 231, std: 212). *N* = 8000 for all panels.

The origin of these oscillations can be traced to the nature of the leading eigenvalues of **Ĵ**. As also reported in [8] and elaborated in Methods, if the leading eigenvalue of **Ĵ**, *λ*_1_, is complex then there is no fixed point solution. Instead we assume in this case that *h* (*t*) undergoes a limit cycle with period *T* and find the solution for *c*_1_ (*t*):

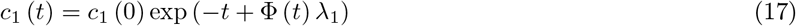

where 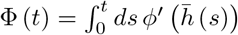.

Requiring that this yield a limit cycle solution for *c*_1_ (*t*) with period *T* yields

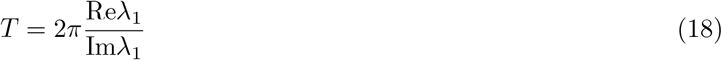

and this additionally requires that the period-average of *ϕ′* (*h* (*t*)) to be equal to the critical value 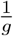.

Indeed, we observe that in the case of a **Ĵ** with complex leading eigenvalue, most realizations exhibit oscillatory fluctuations. The height of the second peak in the autocorrelation, *q*(*τ*), grows gradually with *J*_1_ with a realization-dependent rate (Fig 6B). Furthermore, for most individual realizations the period of the dominant oscillatory peak in the autocorrelation, even within the chaotic regime, is well-predicted by the above theoretical prediction for *T* (Eqn 18, Fig 6C).

For for indivdual realizations, there is a sufficiently large *J*_1_ beyond which the second peak of *q*(*τ*) reaches 1 and the dynamics transition to a pure limit cycle (Fig S3). We note that some realizations with a real leading eigenvalue also exhibit oscillatory components in their chaotic dynamics for certain values of *J*_1_, which we presume relate to complex subleading eigenvalues, but these do not exhibit a transition to pure limit cycle (data not shown).

### Realization-dependence and system-size scaling of the transition out of chaos

We now consider the critical value, 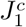, of the strength of structured connectivity that yields a transition out of chaos. In contrast to the case without row balance, we find that in the row-balanced network the transition out of chaos occurs at values of *J*_1_ scaling at least as 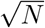. However, the particular value of 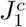 varies considerably across realizations (Fig S2 and S3).

The case of a real leading eigenvalue *λ*_1_ ≈ *g* and the associated transition to fixed point provides a starting point for analyzing the transition out of chaos (a similar argument is made in the Methods for the case of complex leading eigenvalue). In the limit of small residual currents, *δh_i_* ≪ 1, the fixed point equation for the coherent mode current (5) is 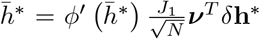. Applying the fixed point requirements derived above that 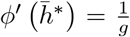 and 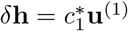, where **u**^(1)^ is the leading eigenvector of **Ĵ**, we find (as also reported in [8]) that

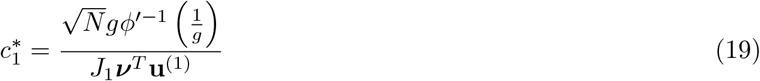

Equation 19 suggests that 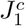 scales as inverse of the overlap between the leading eigenvector, **u**^(1)^, and the output mode, ***ν***. Indeed, we show numerically that the critical value 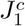 for fixed *g* is negatively correlated with |***ν**^T^***u**^(1)^| (Fig 7A).

**Figure 7.**
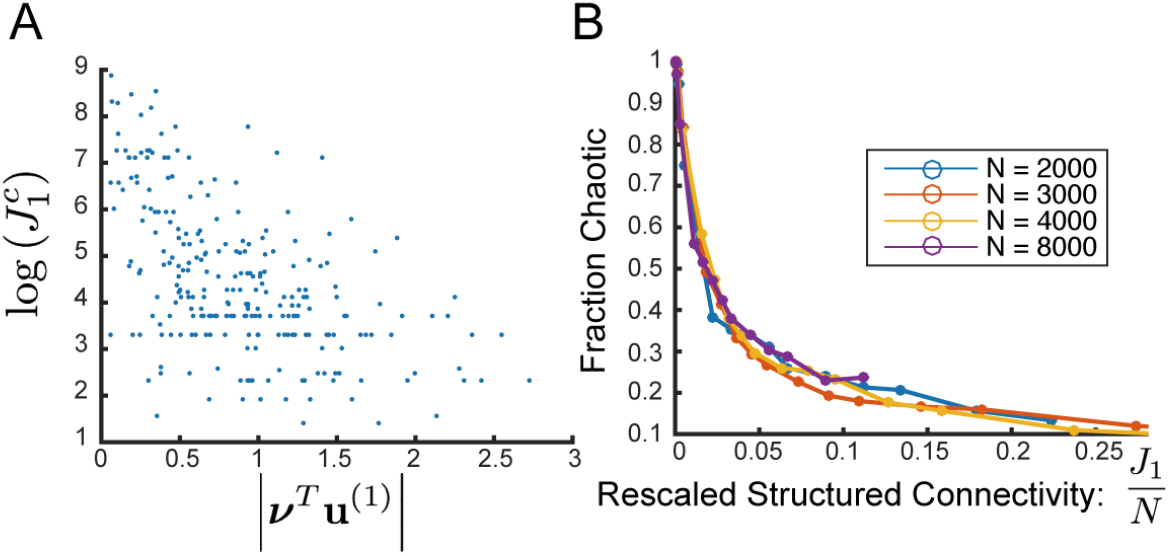
Sufficiently Strong Structure Yields Transition Out of Chaos Despite Row Balance: **(A)** Scatterplot of the logarithm of transitional value of *J*_1_ vs the absolute value of the projection of the output mode, ***ν***, on the leading eigenvector, **u**^(1)^ for 300 connectivity realizations with *N* = 8000. *r*^2^ = 0.29. **(B)** Fraction of realizations displaying chaotic activity as a function of the rescaled structured connectivity: 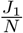. With this scaling the curve appears to be independent of *N*.

We next ask how 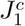 scales with system size. Because the above fixed point assumes 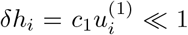 where **u**^(1)^ has norm 1, we must have *c*_1_ be no larger than *O* (1). Since the typical value of |***ν**^T^* **u**^(1)^| is *O* (1) we would naively expect that the typical transition might occur for 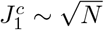. However, we observe numerically that the fraction of realizations exhibiting chaotic dynamics for a given value of *J*_1_ appears to scale as 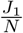 and not as 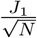 as expected (Fig 7B). For finite *N* chaos appears to be lost for some 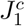 ∼ *N^α^* with *α* ∈ [1/2, 1], and the particular 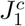 is highly realization-dependent. An analytical derivation of the actual value of 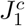 requires a more comprehensive study of the network’s stability.

From our numerical work it appears that in the limit of large system size chaos persists for all values of *J*_1_, for almost all realizations. Indeed for a network with *N* = 16000, for example, we increase *J*_1_ up to values around 100 and observe that chaos persists for most realizations and coexists with a very high degree of spatial coherence. The coherence measure, *χ*, reaches values higher than 0.96 even as fluctuations persist (Fig S4). Thus we conjecture that in the limit *N* → ∞, for almost all realizations, increasing *J*_1_ indefinitely will yield self-tuned coherent chaos with *χ* → 1.

### Multiple Modes of Coherence

We generalize our model in order to construct a network with multiple modes of coherent activity. In this extension we take the structural component, **M**, to be a low-rank matrix comprised of the sum of outer products of pairs of vectors which are all mutually orthogonal:

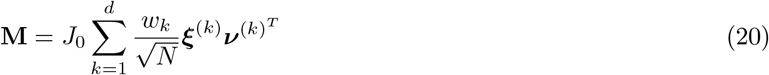

where we have introduced a parameter *J*_0_ that controls the overall strength of the structured connectivity, and a set of parameters *w_k_* satisfying 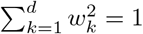 that determine the relative weight of the different modes.

We can extend our schematic representation and think of each row vector 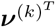 as a separate output mode connected in a feed-forward-like manner to the input mode, ***ξ***^(*k*)^, (Fig 8A) and then decompose the dynamics into the residual dynamics identical to the above (Eqn 4) and the dynamics of the coherent activity along each separate input mode:

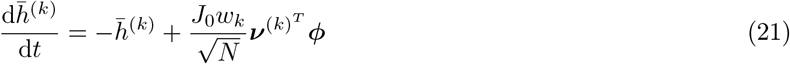

where each separate coherent current is given by 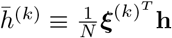.

**Figure 8.**
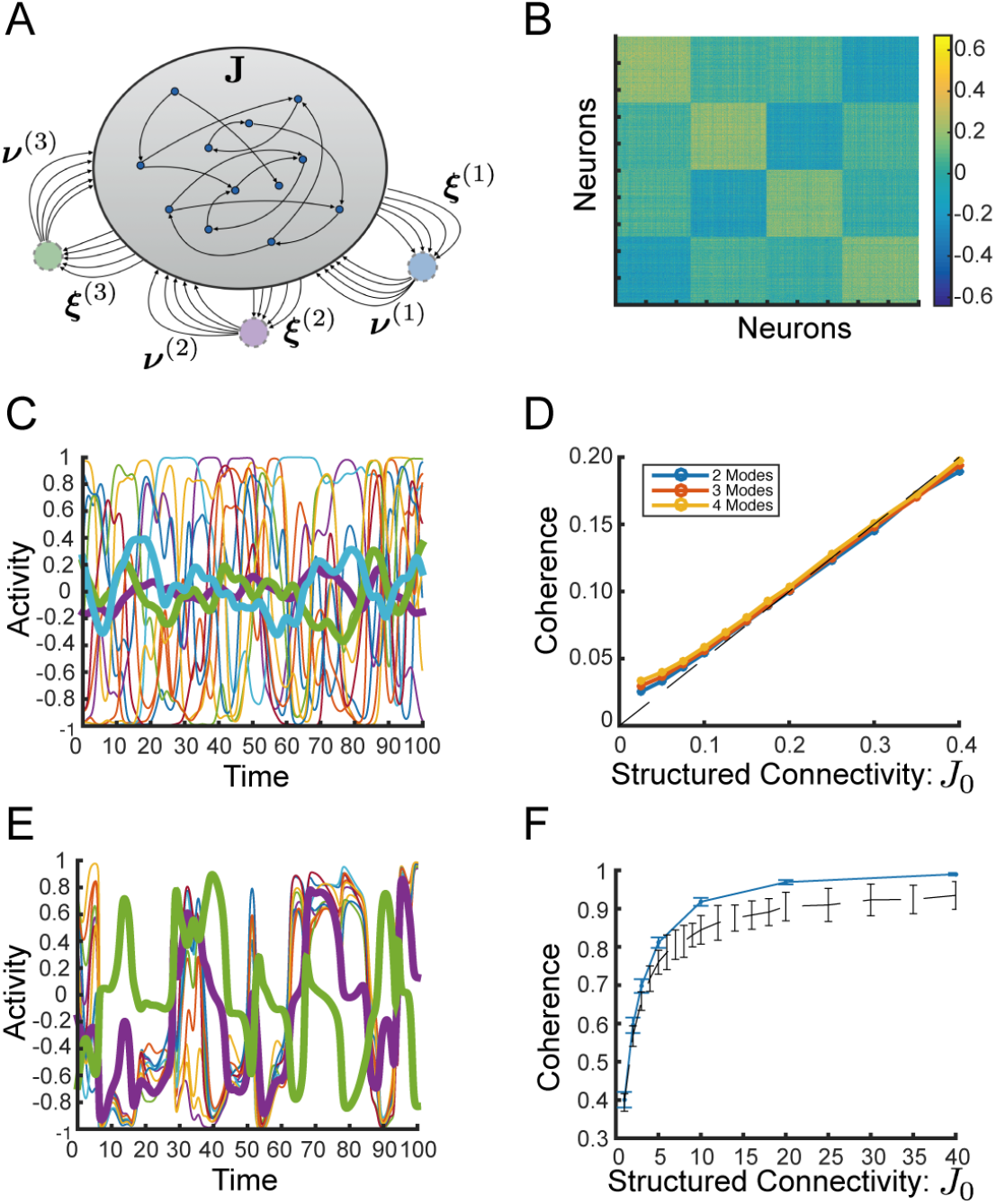
Coherent Chaos Along Multiple Modes. **(A)** Schematic of network with three coherent modes displaying effective output modes, ***ν***^(*k*)^, and input modes, ***ξ***^(*k*)^, each of which are orthogonal to all others. **(B)** Matrix of Pearson correlation coefficients between firing rates, *ϕ_i_*, of pairs of neurons in a network with three coherent modes and *J*_1_ = 1. **(C)** Sample activity trace displaying sample single neuron activities and in thicker lines, three coherent mode activities, *ϕ*^(*k*)^. **(D)** Generalized coherence, *χ*^(*k*)^, as a function of *J*_1_ for 2, 3, 4 modes in the regime of passive coherence. Dashed line displays theory. **(E)** Sample activity traces display extreme coherence for two coherent modes. **(F)** Generalized coherence for two coherent modes (in blue) as a function of *J*_1_ extends to near complete coherence, while chaos persists. Compare network with one coherent mode (in black dashed line). Bar shows standard deviation over 100 realizations. For panels **(B)**, **(C)**, **(E)** *N* = 2000. For panels **(D)** and **(F)** *N* = 4096.

The analytical results found above for the regime of passive coherence can be directly extended to the case of multiple modes. In particular, in the limit where *J*_0_ ≪ 1, the separate coherent modes are independent of each other and driven passively by the residual fluctuations (Fig 8C) such that

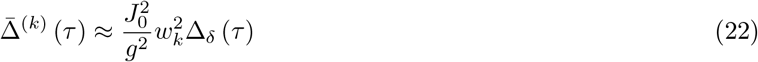

where Δ^(*k*)^ (*τ*) = 〈 *h*^(*k*)^ (*t*) *h*^(*k*)^ (*t* + *τ*)〉 is the autocorrelation function of the *k*th coherent mode.

The resulting covariance matrix, *C_ij_* ≡ 〈*ϕ_i_ϕ_j_*〉, has a low-rank structure which is shaped by the input modes (Fig 8B). In particular, by Taylor expanding *ϕ_i_* around *δh_i_*:

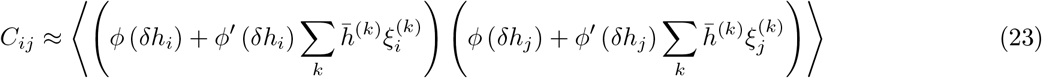

Since the residuals are on average uncorrelated and the distinct input modes are orthogonal, we find using Eqn 22 that to leading order:

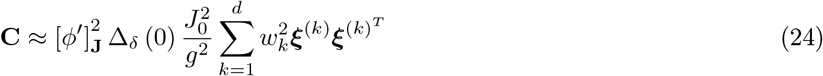

We generalize our measure of coherence to measure the *d*-dimensional coherence, or the fraction of total power which is shared along the *d* input mode directions:

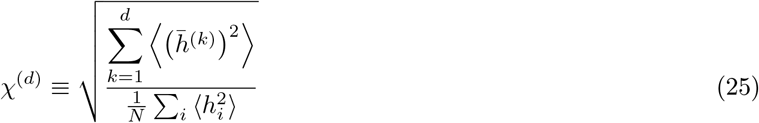

and we find that for *J_k_* ≪ *g* and finite *d*

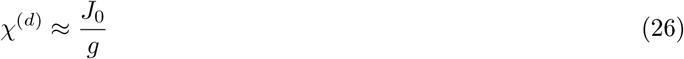

Numerically, this prediction holds well for up to 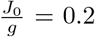 for at least up to *d* = 4 as we show in Fig 8D, and we expect it to hold for larger *d* as well.

We note that in the regime of passive coherence, just as in the case of a single coherent mode, we can relax the restrictions on ***ξ***^(*k*)^ and *ϕ*: Our results hold for ***ξ***^(*k*)^ any norm 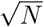 vector orthogonal to ***ν***^(*j*)^ ∀*j* and ***ξ***^(*j*)^ *j* ≠ *k*, and also for *ϕ* any sigmoidal function.

In addition, we can generalize row balance by subtracting the weighted row-average for each input mode such that every ***ξ***^(*k*)^ will reside in the null space of the new connectivity matrix, **J̃**^(*d*)^:

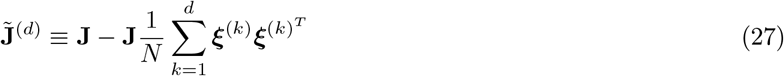

For *d* > 2 this generalized row balance does not appear to preserve chaotic fluctuations and instead fixed points or limit cycles appear for *J*_0_ ∼ *O* (1).

Intriguingly, for *d* = 2 we observe that with generalized row balance the chaotic regime persists as the structured connectivity is strengthened and the dynamics become increasingly coherent. The dynamics display switching-like activity in which at any time one of the two coherent modes appears to be near the critical value *h^c^* while the other mode is near zero (Fig 8E). It appears that the generalized coherence approaches 1 while chaos persists (Fig 8F) such that we conjecture that just as in the case of *d* = 1, here too in the large *N* limit *χ*^(2)^ → 1 as *J*_0_ → *∞*.

## Discussion

Coherent fluctuations are prevalent in cortical activity ranging in spatial scale from shared variability in membrane potential and spiking in local circuits to global signals measured across the scalp via EEG or across voxels via fMRI [40, 27, 1, 14]. Constructing a model that intrinsically generates coherent fluctuations has been a challenge to theorists.

We have studied the intrinsic generation of coherent chaotic dynamics in recurrent neural networks. Our model consists of rate-based neurons whose recurrent connections include a structured component in addition to random connections. The structured component is a low-rank connectivity matrix consisting of outer products between orthogonal pairs of vectors which allow local fluctuations to be summed along an output mode, amplified and projected to an input mode resulting in coherent fluctuations. The orthogonality of input and output mode effectively embeds a purely feedforward structure within the recurrent connectivity, thus avoiding feedback of the coherent fluctuations along the input mode.

In the regime where the structured component is weak, the local synaptic currents are effectively uncoupled from the coherent mode activity and their dynamics are similar to that of a random network with no structured component at all. The local fluctuations are summed by the structured component of connectivity to drive the coherent mode, which follows those fluctuations in a passive manner. In this regime of passive coherent chaos we derive a perturbative dynamical mean-field theory following [30, 12] which shows that the coherence grows linearly with the ratio of the strength of structured connectivity to the random connectivity. We show that this analysis extends to multiple modes of coherent activity yielding a finite-rank covariance pattern for the coherent fluctuations.

For moderate strength of structured connectivity the network exhibits significant realization-dependence and most realizations transition to either a fixed point or a limit cycle. A realization-dependent theory of these transitions is beyond the scope of this work. We add a row-balance constraint, placing the input mode in the null-space of the random connectivity matrix, and we observe that this constraint preserves chaos, reduces the variability between realizations, and increases the level of coherence.

With row balance, increased strength of structured connectivity yields a crossover to a distinct regime of self-tuned coherent chaos. In this regime the network undergoes Up-Down-like switching between two states each of which are characterized by slow, highly coherent dynamics. We show how row balance facilitates this regime by enabling the coherent mode to act as a dynamic gain to the dynamics of the residual currents. Consequently, intermittent marginal dynamics emerge as the coherent mode self-adjusts to one of two critical values. Interestingly the crossover to this self-tuned coherent regime happens for moderate strength of structured connectivity, independently of network size.

In the regime of self-tuned coherent chaos, realization-dependent qualitative differences begin to emerge with increasing strength of structured component, *J*_1_. For realizations of the row-balanced random connectivity with real leading eigenvalue, symmetry-breaking emerges such that individual initial conditions yield trajectories that spend more time near one of the critical values of the coherent mode than the other. For realizations with complex leading eigenvalue, oscillatory fluctuations begin to emerge. The frequency of these oscillations is well predicted by the phase of the leading eigenvalue. Note that we have not addressed the question of the necessary scaling of *J*_1_ for the emergence of realization-dependence in the chaotic regime for the limit of large system size.

As structured connectivity is further strengthened chaos persists even as coherence continues to increase until the dynamics are dominated almost entirely by the one dimensional fluctuations of the coherent mode. For a finite network, above some critical strength of the structured component the system converges to either a fixed point or a limit cycle, depending on the leading eigenvalue of the row-balanced random connectivity. Our numerical work suggests that the critical strength of structured connectivity grows with the system size, most likely scaling as 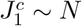 (at least as 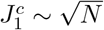). Hence we conjecture that for the scaling of the strength of structured connectivity presented here, as the network size diverges coherent chaos persists independent of *J*_1_ for most realizations, and the level of coherence can be brought arbitrarily close to 1.

Importantly, in the regime of weak structured connectivity and passive coherence some of the assumptions of our model can be loosened. First, in this regime row balance on the random connectivity is not necessary. Additionally, we need not require the input mode be binary but rather any general pair of orthogonal vectors can serve as input and output mode. Moreover we can loosen the restriction on orthogonality: a random pair of vectors can be used without qualitative impact on the dynamics presented here because the contribution of the realization-dependent overlap between the two vectors in this regime will be negligible relative to the typical contribution from the full-rank random connectivity, **J**.

On the other hand, achieving self-tuned and highly coherent chaos requires the network be finely-tuned to a high degree. The orthogonality of the input and output modes is not enough in order to achieve highly coherent chaos because of interactions between the random and structured components of connectivity. We therefore constrain the random component to satisfy row balance by ensuring that the input mode of the structured connectivity be in the null-space of the random component of connectivity. In addition to row balance, the self-tuned coherence regime depends on the choice of the single-neuron transfer function and input-mode vector. In the case where the transfer function is an odd function, such as tanh used throughout the main text, the input mode can be any binary vector. Otherwise, high coherence is achieved only for a uniform input mode, *ξ_i_* = 1 ∀*i*. In the latter case, the theory developed here predicts coherent fluctuations that switch between two non-symmetric values of the coherent mode, corresponding to the two points where where the slope of the transfer function equals 1/*g*, and we have verified this numerically (Fig S1).

An interesting question is whether the particular structure of the connectivity matrix in our model can be achieved by a biologically plausible synaptic learning rule. Prior studies of sequence generation have constructed learning rules that yield connectivity which is comprised of outer-products of random vectors [29, 33] and these could form the basis for learning the necessary orthogonal rank-one structure. Plausible learning rules for yielding balanced excitation-inhibition dynamics [39, 15, 13] could potentially provide a foundation for learning row balance. It is thus plausible that the constraints of our model can be achieved by an appropriate synaptic learning, especially for the more robust regime of mild coherence. On the other hand, it is unclear to us whether the high degree of fine tuning required for the self-tuned coherence regime can be achieved by a biologically plausible learning rule. Investigating candidates of appropriate learning rules for generating coherent chaos, is beyond the scope of this work.

In the case of a uniform input mode the model can be constructed as an excitation-inhibition network, for example with half the neurons defined as excitatory by setting *ν_i_* = 1 and the other half defined as inhibitory via *ν_i_* = −1 (or a larger inhibitory value to compensate for a smaller fraction of inhibitory neurons). From this perspective the coherent fluctuations, in particular in the regime of passive coherence, can be understood within the framework of dynamic excitation-inhibition balance [38]. In this case the pair of balance equations are degenerate and constrain only the mean excitatory population rate to be nearly equal the inhibitory rate, but otherwise leave the overall mean rate unconstrained. Local residual fluctuations yield only small differences in mean population rates, thus leaving the balance satisfied, but these small differences drive significant coherent fluctuations because of the strong balanced connectivity. In the general setting of excitation-inhibition balance the pair of balance equations fully determine the mean rates to leading order and no coherent fluctuations are possible without introducing shared fluctuations in the external drive. We note that excitation-inhibition networks in the literature have sometimes been constructed yielding degenerate balance equations. As we have shown here, such choices have dramatic impact on the dynamics and the results should not be assumed to be generalizable.

In parallel to our study a pre-print has been published which explores a very similar model[10]. The authors observe a similar phenomenon as the self-tuned coherence studied here, and attempt to explain it by an iterative numerical solution of locally Gaussian dynamic mean-field equations. They do not make the role of row balance clear in their analysis. In contrast, we have focused on analytical solutions in the limit of both weak and strong structured connectivity, deriving a perturbative dynamic mean-field solution for the regime of weak structure and passive coherence. As we have shown here row balance is critical for moderate structure and self-tuned coherent chaos. Additionally we have shown that a full understanding of the properties of the highly coherent regime requires a realization-dependent mean-field analysis. In particular, we have explained that the leading eigenvalue of the row-balanced random connectivity matrix impacts qualitative features of the chaotic dynamics, yielding either broken symmetry or oscillatory fluctuations. Furthermore the critical strength of structured connectivity that leads to a transition to either fixed point or limit cycle is correlated with the extent of overlap between the leading eigenvector and the output mode.

As mentioned in the main text and introduction, a previous study also explored the case of a single orthogonal E-I structured component[8]. They derived the fixed point and limit cycle solutions which we reviewed here, but did not focus on the chaotic regime and they did not discuss the role of network size in the transition out of chaos. Our focus here was on the chaotic regime, both the emergence of coherence for small structured connectivity and the imprint of the non-chaotic regime on the chaotic dynamics for moderate structured connectivity.

A separate study has claimed to observe coherent activity in excitation-inhibition networks of spiking neurons [37]. A study of the dynamics of spiking neurons is beyond the scope of our work, although we would conjecture that coherent activity would arise with orthogonal, rank-one E-I structure in that setting as well.

Previous work has shown how shared inputs from external drive can drive correlated fluctuations in excitation-inhibition networks [6, 25]. In our current work, in the context of rate neurons, we show that such correlated fluctuations can be generated internally by a recurrent network without external drive. In order to avoid either saturation or pure oscillations the coherent activity mode must not drive itself through a feedback loop. In order to achieve this it is necessary that the structured component embed an effectively-feedforward projection between a pair of orthogonal modes. In parallel, Darshan et al [5] have developed a theory for internally generated correlations in excitation-inhibition networks of binary units. The underlying principle is similar: the recurrent connectivity embeds a purely feedforward structure.

We note that the structured component of connectivity in our network is non-normal. The dynamics of non-normal matrices have drawn a fair amount of interest with suggested functional impact on working memory [7, 9] and amplification [17]. Non-normal matrices embed feedforward structure within recurrent connectivity, and the resulting dynamics even in a linear system are not fully determined by the eigenspectrum but depend on the structure of the corresponding eigenvectors[36]. It has been shown that E-I networks are generally non-normal, and that rank one E-I structure amplifies small differences between excitatory and inhibitory rates driving a large common response [17, 11]. This amplification is related to the way in our network, small fluctuations of the residuals are summed along the output mode and drive coherent fluctuations along the input mode in the regime of passive coherence, but these fluctuations are internally driven by the non-linear dynamics whereas the dynamics in those previous studies were linear. As pointed out in [2], a structured component such as in our model is purely feedforward and can be considered an extreme case of non-normality as it has only zero eigenvalues and therefore all the power in its Schur decomposition is in the off-diagonal. The results here depend on this property and cannot be extended to connectivity with only partial feedforward structure.

Rate model dynamics with a rank-one structured component have been studied in depth recently [24, 16]. Since these works focused on time-averaged activity and not fluctuations they did not observe coherent activity in the case of an outer product of a pair of orthogonal vectors as studied here. These works also differed in that the strength of the structured connectivity was scaled as 1/*N*. This scaling is similar to our limit of weak structured connectivity, *J*_1_ ≪ 1, and guarantees that dynamic mean-field theory holds in the limit of large system size, but in that scaling coherent fluctuations will appear only as a finite-size correction.

It has been previously observed [22] and then proven [34] that adding an orthogonal outer-product to a random matrix generates realization-dependent outliers in the eigenspectrum, and furthermore that these outliers are be removed by row balance. It has been previously observed that performing such a subtraction has significant impact on the resulting dynamics [8, 31]. Yet the relationship between the change in eigenspectrum and the dynamics has not been made clear beyond the basic observations regarding the stability of a fixed-point at zero. Here we suggest that the impact of row balance on the chaotic dynamics is not directly related to the eigenspectrum but that this adjustment should be thought of as effectively subtracting the coherent-mode activity from each individual neuron, thus -preventing feedback loops to the coherent mode. We show that row balance enables the emergence of slow residual dynamics with the coherent mode playing the role of dynamic gain, and that it is crucial for the emergence of self-tuned chaos and highly coherent dynamics.

In conclusion we have presented a simple model which generates coherent chaos in which macroscopic fluctuations emerge through the interplay of random connectivity and a structured component that embeds a feedforward connection from an output mode to an orthogonal input mode.

## Methods

### Exact Decomposed Dynamics and Row Balance

We write the full dynamics without row balance:

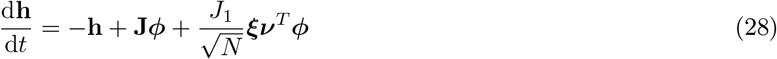

and we define

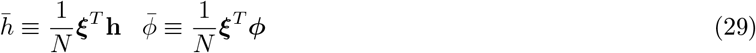

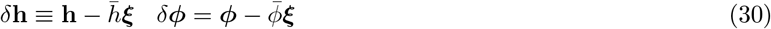

Applying these definitions to the full dynamics (and noting that ***ν**^T^**ϕ*** = ***ν**^T^δ**ϕ***), the exact coherent mode dynamics are

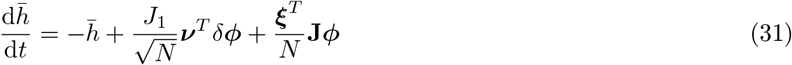

and by subtracting these from the full dynamics of **h**, the decomposed dynamics are:

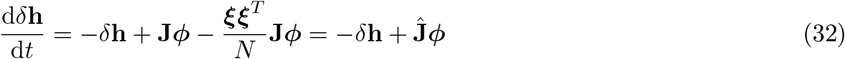

where 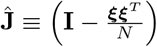 **J** as introduced in the main text.

We observe that the constraint ***ξ**^T^ δ***h** = 0 must be satisfied automatically by the residual dynamics (Eqn 32), and this can be confirmed by verifying that

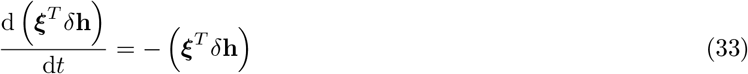

In the regime where *J*_1_ ≪ 1 the *ϕ_j_* are nearly uncorrelated and therefore 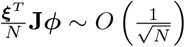 and can be ignored.

In the regime with strong structured connectivity we must consider this term in Eqn 31. To that end we write ***ϕ*** = *ϕ****ξ*** + *δ**ϕ*** and also write the transformation of the input mode via the random matrix as

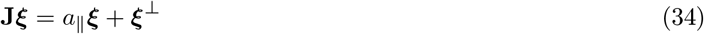

where *a*_||_ is a realization-dependent scalar and ***ξ***^⊥^ is a realization-dependent vector orthogonal to ***ξ***. That yields coherent mode dynamics

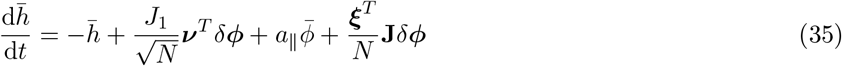

and residual dynamics

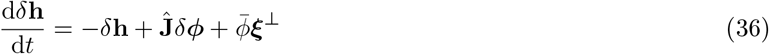

As *J*_1_ increases and the fluctuations in the coherent activity, *ϕ* (*t*), drive feedback in two ways. First of all, **J** maps the coherent activity back along the input mode driving direct feedback to the coherent current *h* via the term *a*_||_ *ϕ*.

Secondly, **J** maps the coherent activity in a realization-dependent direction, ***ξ***^⊥^, orthogonal to the input mode. This drives the residual activity fluctuations *δ***h** via the term *ϕ****ξ***^⊥^, and this biasing of the residual fluctuations may in turn generate feedback to the coherent current through the output mode via ***ν**^T^ δ**ϕ***.

Both of these feedback terms are realization dependent, and both of them are canceled via the row balance sub-traction

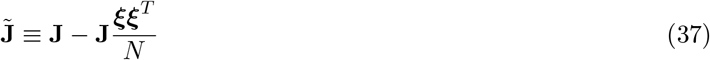

which yields exact coherent mode dynamics

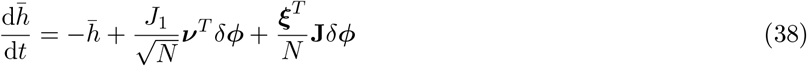

and residual dynamics

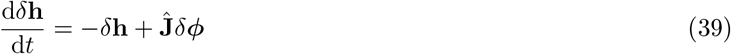

And in this case the residual dynamics are again uncorrelated so that 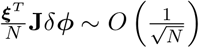 and can again be ignored in the coherent mode dynamics.

Note that *ϕ* no longer drives feedback to either the residual or the coherent dynamics. Nevertheless the dynamics are still coupled in both directions as *δ**ϕ*** depends on *h*.

### Perturbative Dynamic Mean-Field Theory in the Limit of Weak Structured Connectivity

We derive the dynamic mean-field equations in the limit of small *J*_1_ using a perturbative approach. We write the mean-field dynamics of the residuals as

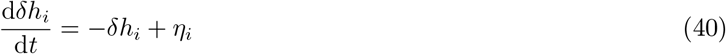

and the coherent component as

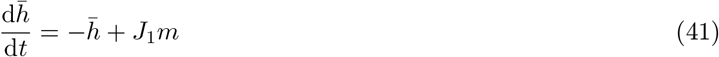

where *η_i_* and *m* are assumed to be uncorrelated, mean-zero Gaussians. For general *J*_1_ the assumption of Gaussianity fails, therefore we assume *J*_1_ ≪ 1.

The autocorrelation of *η_i_* is given by

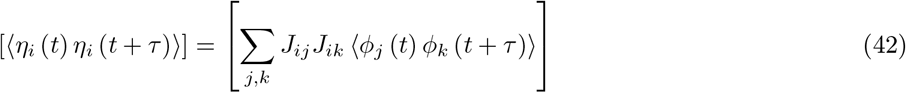

where we have introduced [] as the notation for averaging over realizations. We assume that *ϕ_j_* is independent of *J_ij_* and so the terms *j* ≠ *k* have average zero over realizations and we get

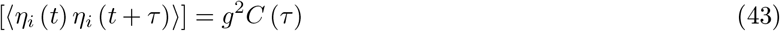

where

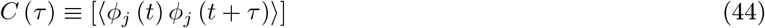

The autocorrelation of *m* is given by

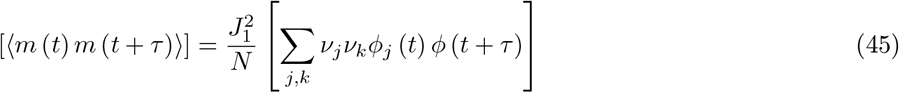

And again the *j* ≠ *k* terms fall in the realization average so that

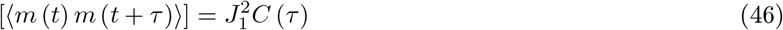

Next we define the autocorrelation of the residuals

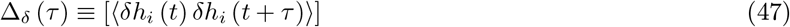

and the autocorrelation of the coherent current

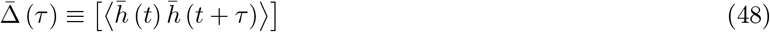

and we can follow previous work [30, 12] and write the dynamic mean-field equations for Δ_*δ*_ (*τ*) as

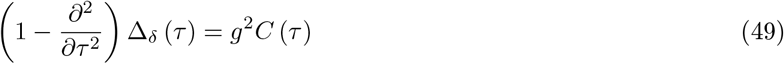

and for Δ(*τ*) as

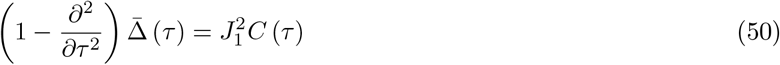

Next we note that for *J*_1_ ≪ *g* we assume that |*h*| ≪ 1 so we have *ϕ_i_* = *ϕ* (*δh_i_* + *ξ_i_h*) ≈ *ϕ* (*δh_i_*) + *ξ_i_ϕ′*(*δh_i_*) *h*. Therefore we have that to leading order

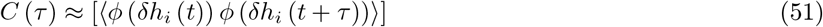

and then following previous results [30, 12] we can write this to leading order as an integral over Gaussians:

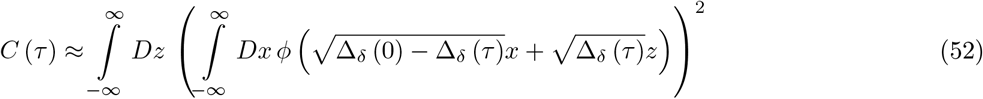

Note that it is possible to compute the sub-leading correction term as well, but for our purposes this is unnecessary. We suffice it to observe that to leading order, the self-consistency equation for Δ_*δ*_ (*τ*) (Eqn 49) reduces to the identical equation for that of a random network without structured component (*J*_1_ = 0)[30], and that Δ (*τ*) contributes only to the sub-leading correction. Following [30, 12] then Eqn 49 can be solved yielding Δ_*δ*_ (*τ*) ≈ Δ_0_ (*τ*), where Δ_0_ (*τ*) is the autocorrelation when *J*_1_ = 0.

The dynamic equation for Δ (*τ*) is identical to that for Δ_*δ*_ (*τ*) except with *J*_1_ in place of *g*, so we conclude that the resulting leading order autocorrelation of the coherent mode is

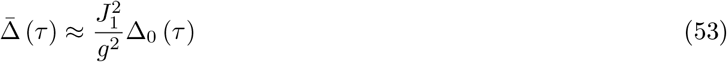

Thus for *J*_1_ ≪ *g* fluctuations in the coherent input are driven passively by the random source which is generated self-consistently by the residual fluctuations, and the resulting autocorrelation of the coherent mode is simply a scaled version of the autocorrelation of the residuals.

It is worth noting that for *J*_1_ ∼ *g* the assumption of Gaussianity is broken due to the cross-correlations between the *ϕ_j_*.

### Analysis of Fixed Points and Limit Cycles in the Limit of Strong Structured Connectivity

In the limit of large *J*_1_ we assume *δh_i_* ≪ 1, and approximate *ϕ_j_* ≈ *ϕ* (*ξ_j_h*) + *ϕ′*(*h*) *δh_j_*, where we have made use of the symmetry of the transfer function and the binary restriction on *ξ_j_*. Note that this linearization clearly holds without symmetric transfer function for the case of uniform *ξ_j_* = 1 as well.

Following the exact decomposition above (Eqns 32 and 31) this yields dynamical equations:

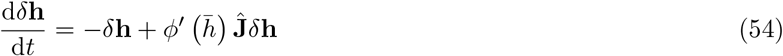

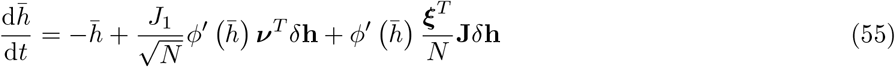

Note that the leading term of the linearization, *ϕ* (*ξ_i_h* = *ξ_i_ϕ* (*h*) has fallen from all terms, so that in this regime the coupling between residuals and coherent mode has been greatly simplified: The residuals serve as the input while the coherent mode acts as a dynamic gain. This reflects the important role of the orthogonality of ***ν*** and ***ξ*** on the one hand, and the row balance subtraction we made to the random connectivity, both of which ensure that the coherent mode does not drive a feedback loop.

In this regime *h* acts as a dynamic gain on the local synaptic currents through *ϕ′* (*h*). Given *h* the equation for the residual currents is linear and therefore their dynamics can be decomposed in the eigenbasis of the matrix

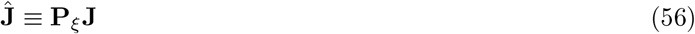

where 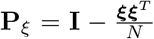 is the projection matrix onto the the subspace orthogonal to ***ξ***. We observe a fine-point not noted in [8]: Had we ignored the constraint ***ξ**^T^ δ***h** = 0 then the residual dynamics would have been determined by **J** and its eigenvalues, and these are not the same as those of **Ĵ**.

We claim that **Ĵ**= **P**_*ξ*_**J** and **J**̃ = **JP**_*ξ*_ have the same eigenvalues. If *λ* is an eigenvalue **J** such that the associated eigenvector **u** is orthogonal to ***ξ***, then clearly **u** is also an eigenvector of both **Ĵ** and **J**̃ with the same eigenvalue *λ*. Otherwise *λ* is an eigenvalue of both **Ĵ** and **J**̃ if and only if (**J** − *λ***I**)^*−*1^ ***ξ*** = **u** is orthogonal to ***ξ***. In this case it can be readily verified that **u** is an eigenvector of **Ĵ**, and **Ju** = *λ***u** + ***ξ*** is an eigenvector of **J**̃. Thus we note also that the eigenvectors of **Ĵ** are orthogonal to ***ξ*** (except potentially for one eigenvector with zero eigenvalue).

We write the eigenvectors as **u**^(*i*)^ with **Ĵ** *·* **u**^(*i*)^ = *λ_i_***u**^(*i*)^. Writing the vector of residual current as 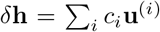 yields dynamics

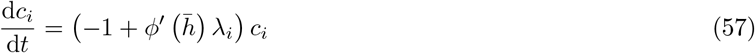

The only (marginally) stable, non-zero fixed point is achieved with *c*_1_ ≠ 0 and *c_i_* = 0 for all *i >* 1. And the fixed-point equation is

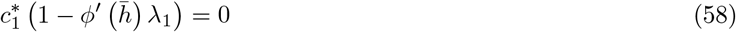

This fixed point only exists if *λ*_1_ is real, and yields a fixed-point requirement for *h*∗:

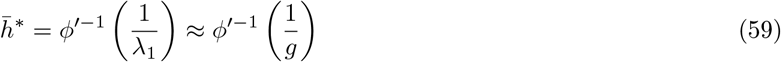

In order to close the loop we turn to the fixed point equation for the coherent dynamics. Ignoring the term 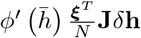, which yields an 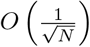 correction we find:

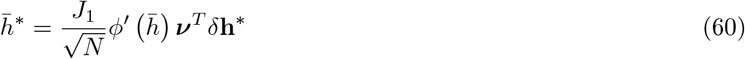

which in turn, using *δ***h**^∗^ = *c*_1_**u**^(1)^, yields a solution to leading order for *c*_1_: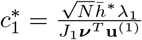, as reported in [8].

If *λ*_1_ is complex there is no fixed point but rather a limit-cycle solution to the dynamics of the complex-valued *c*_1_ exists with *δ***h** (*t*) = Re *c*_1_ (*t*) **u**^(1)^, and *c_i_* = 0 for all other eigenmodes. Assuming *h* (*t*) is periodic with period *T*, we can separate variables and integrate Eqn 57 in order to find *c*_1_ (*t*) is given by

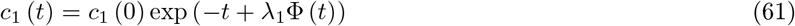

for *t* ≤ *T*, where 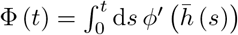. Writing *c*_1_ (0) = 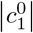 exp (*iθ*_0_) and using Re*λ*_1_ ≈ *g*, this gives

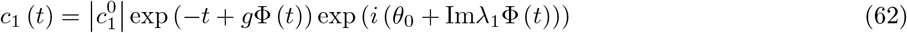

A limit cycle in phase with *h* (*t*) means *c*_1_ (*T*) = *c*_1_ (0) and this requires that both *g*Φ (*T*) = *T* and also Im [*λ*_1_] Φ (*T*) = 2*π*. From the first requirement we find that the average value of *ϕ′*over a period must be the critical value:

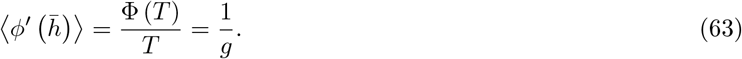

And combining the second requirement yields an expression for the period (Eqn 18, as reported in [8] as well):

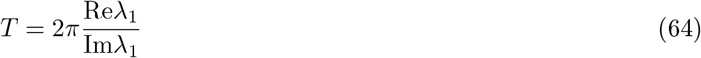

We can further write a self-consistency expression for *h* (*t*) by taking *c*_1_ (*t*) as given by Eqn 62 and integrate over the coherent mode dynamics:

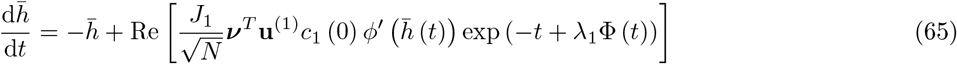

which yields

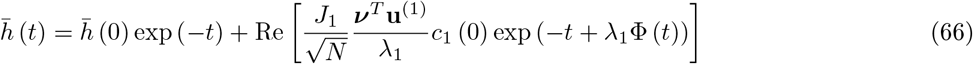

Without loss of generality we assume that *h* (0) = *h^c^* = *ϕ*^′−1^ and 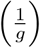, then

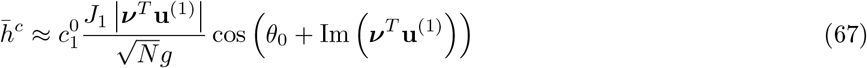

This is analogous to the fixed point equation for *h*^∗^ and 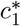. In both cases the requirement that 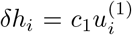 ≪ 1 requires that *c*_1_ be maximally *O* (1) and motivates our conjectures about the realization-dependence and system-size scaling of the transition out of chaos. In particular, we expect and confirm numerically that the critical value of *J*_1_ for transition to either fixed point or limit cycle is inversely proportional to |***ν**^T^* **u**^(1)^| and grows with network size (see main text and Fig 7).

In the case of complex leading eigenvalue, simulations confirm that a projection of the full synaptic current dynamics into the coherent mode and the leading eigenvector plane (consisting of real and imaginary parts of **u**^(1)^) accounts for well over 0.99 of the total variance. Even restricting ourselves to the variance of the residuals, *δh_i_*, we find that 0.98 of the variance is restricted to the leading eigenvector plane (Fig S3).

For *N* = 4000 we simulate 219 realizations of random connectivity with complex leading eigenvalue and find that for sufficiently large *J*_1_ all but one of these realizations yield highly oscillatory dynamics with period predicted nearly perfectly by theory (Fig S3).

We note that in the limit of large *N* we expect that the typical size of the imaginary component of the leading eigen-value, *λ*_1_, shrinks such that the typical period grows. These longer period oscillations are characterized by square-wave-like shape in which the dynamics of the coherent component slows around the critical value 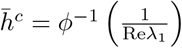, which is identical to the fixed-point value of *h* when *λ*_1_ is real. (Fig S3)

The fraction of realizations with real leading eigenvalue in the large *N* limit has not been calculated analytically to our knowledge. We find numerically that this fraction appears to saturate roughly around 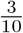 for *N* ≳ 8000.

### Lyapunov Exponent, Limit Cycles, and Fixed Points

In order to calculate the largest Lyapunov exponent, we begin with a point **h**_0_ along the trajectory of the dynamics and we solve concurrently for the dynamics of the trajectory, **h** (*t*) with **h** (0) = **h**_0_, and for a randomly chosen perturbation, ***η*** (*t*). The trajectory **h** (*t*) yields the time-dependent Jacobian matrix for each point along the trajectory:

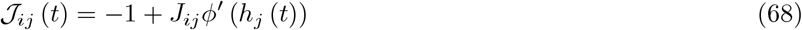

We choose a random unit-norm vector ***η*** (0) = ***η***_0_ and iterate the linearized dynamics of the perturbation:

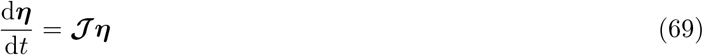

The largest Lyapunov exponent is given by

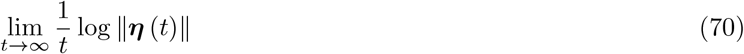

In practice we iterate 69 until *t* = 5000, that is, 5000 times the intrinsic time-scale of the dynamics, and we renormalize ***η*** (*t*) at intervals of *t* = 100*n* for *n* = {1, 2, …, 50}.

We classify fixed points numerically by a threshold on the fluctuations of the coherent input: std (*h* (*t*)) *≤* 5×10^−4^.

We classify limit cycles by a threshold on the second peak of the normalized coherent autocorrelation: *q_peak_ ≥* 0.9.

We confirm that all of the trials with negative Lyapunov exponent were categorized as either fixed points or limit cycles. A small fraction of trials classified as limit cycles had positive Lyapunov exponents but with the largest one 0.0043.

## Acknowledgements

We thank Rainer Engelken for advice on numerically calculating Lyapunov exponents, and Yu Hu for very helpful comments on the manuscript. We thank the anonymous reviewers for thorough questions that helped clarify the manuscript.

## Supporting Information Legends

**Figure S1 (see Fig 4). Self-Tuned Coherent Dynamics With Non-Symmetric Transfer Function. (A)** We use a non-symmetric transfer function *ϕ* (*h*) = (1 + exp *−βh*)^*−p*^ with *β* = 4 and 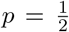. **(B)** Activity trace of coherent activity *ϕ*(*t*) in black and 10 randomly chosen neurons *ϕ_i_* (*t*) displays coherent switching between slow states. **(C)** Histogram of values of coherent current, *h*, displays bimodality with peaks near the critical values predicted by theory where *ϕ*′ (*h*) = 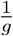. Simulations for *N* = 2000 and *g* = 2 with row balance.

**Figure S2 (see Fig 5). Real Leading Eigenvalues Yield Fixed Points. (A)** Sample chaotic dynamics for *J*_1_ = 0.1. **(B)** Sample dynamics of same connectivity realization as in (A) but with *J*_1_ = 1. **(C)** Scatterplot of all 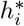 at fixed point, plotted against individual components of the leading eigenvector, 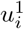. Red dot is value of coherent mode at FP. Black dashed line is FP value predicted from theory. **(D)** Value of coherent mode at FP, *h*^*^, as a function of the standard deviation of the random connectivity, *g*. Black line is prediction from theory: *h^c^* 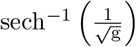 **(E)** Phase diagram for a single realization with real leading eigenvalue. Colormap shows the absolute value of the mean coherent current over a single trial, |〈 *h*〉| which is close to zero when the network is chaotic and non-zero when at a fixed point. The bar below shows the fixed point value predicted by theory, *h*^*c*^, which is independent of *J*_1_. **(F)** Stability eigenvalue at fixed point, i.e. leading eigenvalue of the Jacobian, 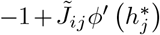, as a function of *J*_1_ for a specific realization of the random connectivity. The fixed point exhibits marginal stability independent of *J*_1_. Networks in all panels have row balance. *N* = 4000.

**Figure S3 (see Fig 6). Complex Leading Eigenvalues Yield Limit Cycles. (A)** Top: Sample chaotic dynamics for *J*_1_ = 0.25. Bottom: Autocorrelation of coherent mode shows oscillatory ringing. **(B)** Same connectivity realization as (A) but with *J*_1_ = 2. Dashed pink line in top panel is prediction from solving the three-dimensional dynamics. Autocorrelation shows near-perfect oscillations. **(C)** Projection of the full dynamics of (B) onto coherent mode and the real and imaginary parts of the leading eigenvector. These three dimensions account for more thanof the total variance of the dynamics. Gray projection onto the leading eigenvector plane accounts for 0.98 of the variance of the residual currents. **(D)** Scatterplot of period of oscillations plotted against the phase of the leading eigenvalue, 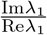, of **J**̃, for 219 different realizations of the random connectivity. Black line shows prediction from theory, *T* = 2π 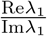. **(E)** Phase diagram for a single connectivity realization with complex leading eigenvalue. Colormap shows the second peak of the normalized autocorrelation of the coherent mode. Networks in all panels have row balance. *N* = 4000

**Figure S4 (see Fig 3). Near Perfect Coherence as Strength of Structured Connectivity is Increased. (A)** Plot of coherence, *χ*, vs strength of structured connectivity, *J*_1_, for networks of size *N* = 16000 with row balance. Dots display average over realizations, bars display standard deviation. Only chaotic realizations included (those not found to be at a fixed point or a limit cycle - see methods). More than 20 realizations per value of *J*_1_. For *J*_1_ = 100, 22 out of 30 realizations were chaotic and the average coherence among these realizations was 0.963.

